# Progressive matrix stiffening of tyramine-modified silk fibroin hydrogels governs stage-specific pulmonary fibroblast activation

**DOI:** 10.64898/2026.06.01.729382

**Authors:** Mariah L. Arral, Maria Savvidou, Adam S. Mullis, Aria Z. Yang, Thomas Falcucci, Julie Leonard-Duke, Pamela L. Graney, Sabrina Madiedo-Podvrsan, Sanjana Gopalakrishnan, Jugal Kishore Sahoo, Jing-Jie Huang, Gordana Vunjak-Novakovic, David L. Kaplan

## Abstract

Fibrosis is a progressive and often fatal pathological process characterized by excessive extracellular matrix deposition, tissue stiffening, and irreversible organ dysfunction. Effective antifibrotic therapies remain limited by the lack of in vitro models that recapitulate the full spectrum of fibrotic disease progression. Here, we leverage tyramine-modified silk fibroin (SF-TA) hydrogels to investigate normal human lung fibroblasts (NHLF) responses to progressively stiffening environments relevant to pulmonary fibrosis. Two hydrogel formulations with distinct stiffening profiles over 14 days were prepared: a gradual-stiffening 0% SF-TA formulation reaching ∼20 kPa, and a rapidly stiffening 50% SF-TA formulation reaching ∼60 kPa. NHLFs were cultured on both formulations, with and without TGFβ (5 ng/mL), for 14 days and assessed for viability, metabolic activity, cytokine and collagen secretion, cytoskeletal organization, and mechanotransductive gene expression. The 0% SF-TA hydrogels drove sustained fibroblast proliferation and elevated secretion of IL-6, IL-8, and MCP-1, consistent with early inflammatory fibrosis. The 50% SF-TA hydrogels induced a metabolic plateau without senescence, suppressed inflammatory cytokine secretion, and, in the presence of TGFβ, led to significant upregulation of ACTA2 and CTGF, alongside α-SMA stress fiber incorporation, consistent with established myofibroblast persistence. Both conditions produced comparable secreted collagen output by day 14. Together, these findings establish dynamically stiffening SF-TA hydrogels as a tunable platform for investigating stage-dependent fibroblast activation and mechanobiological progression in fibrosis.

## Introduction

Fibrosis is a progressive and often fatal pathological process characterized by excessive extracellular matrix (ECM) deposition by myofibroblasts, inflammation, tissue stiffening, and eventually irreversible loss of organ function[1,2]. Fibrotic diseases are estimated to contribute to nearly 35% of all global deaths and can affect virtually any organ [2–4]. Pulmonary, hepatic, and cardiac manifestations are among the greatest disease burdens in terms of mortality[2,5]. Despite its prevalence across tissues and organ systems, effective antifibrotic therapies remain limited, reflecting the complex, multifactorial nature of fibrotic disease progression[5]. The lack of antifibrotic treatments directly results in the only curative option for late-stage disease being organ transplantation[5].

Pulmonary fibrosis is characterized by progressive scarring of the lung parenchyma, leading to impaired gas exchange and respiratory failure[6]. Idiopathic pulmonary fibrosis, the most severe form, is incurable and fatal, with a median survival of only 2 to 5 years[7,8]. The progression of pulmonary fibrosis begins with the accumulation of fibroblasts and myofibroblasts within stiffened lung tissue, which promotes excessive matrix deposition and architectural distortion[3,9,10]. Healthy lung parenchyma exhibits an elastic modulus of ∼2 kPa[11–13], whereas fibrotic tissue can range 10-100 kPa[14,15]. This mechanical transformation reinforces profibrotic signaling through cytoskeletal tension, focal adhesion remodeling, and integrin-mediated matrix sensing[5,9,16]. In parallel, profibrotic cytokines, particularly transforming growth factor-beta (TGFβ), drive fibroblast differentiation into contractile, matrix-producing myofibroblasts [17,18]. These myofibroblasts are characterized by the incorporation of alpha-smooth muscle actin (α-SMA) into stress fibers that are resistant to apoptotic resolution[16,19]. Together, these mechanical and biochemical cues create a self-reinforcing cycle of myofibroblast activation that drives progressive fibrotic disease.

In vitro models of fibrosis have sought to recapitulate the mechanical and biochemical features of fibrotic tissue by incorporating native ECM components, such as collagen[20–22] and, more recently, decellularized matrices[3,23,24]. These systems have yielded important mechanistic insights; however, decellularized matrices rely on materials derived from animal or patient tissue, which limits availability and standardization. Collagen-based systems, while accessible and widely used, pose challenges for long-term culturing required to mimic the fibrotic environment[20,25]. Furthermore, most existing models present static mechanical environments and fail to capture the progressive, time-dependent stiffening that characterizes fibrotic disease evolution in vivo [16,26–29]. The concept of mechanical memory further suggests that the trajectory of stiffening, not just the endpoint modulus, may be a critical determinant of fibroblast fate in progressive fibrotic disease[30,31]. There remains a need for a chemically defined hydrogel platform that enables systematic, controlled investigation of mechanical cues and profibrotic signaling in fibroblast activation.

Silk fibroin-based hydrogels offer a well-defined platform to address the limitations of existing fibrosis models[32–35]. Derived from *Bombyx mori* silkworms, silk fibroin is a commercially available non-mammalian protein polymer that does not require vertebrate animal sacrifice or patient tissue procurement[36]. This avoids logistical challenges associated with collagen or decellularized matrix systems. Additionally, chemical modification of silk fibroin enables precise tuning of mechanical properties over physiologically relevant ranges [10,33,36]. Tyramine modification, in particular, enables horseradish peroxidase-mediated crosslinking, yielding hydrogels with controllable, temporally tunable stiffness. This allows hydrogel stiffness to evolve in a controlled manner that mimics the progressive mechanical changes in fibrosis[33].

In this study, we leveraged tyramine-modified silk hydrogels to investigate normal human lung fibroblast (NHLF) responses to mechanically dynamic environments relevant to pulmonary fibrosis. Using two gel formulations with distinct stiffening profiles, with or without TGFβ supplementation, we examined fibroblast viability, metabolic activity, inflammatory cytokine and collagen secretion, cytoskeletal organization, and mechanotransductive gene expression over a 14-day culture period. Our findings reveal that distinct mechanical environments differentially drive inflammatory fibroblast activation and persistent myofibroblast differentiation, recapitulating features of both early and established stages of fibrotic disease. Collectively, this work establishes tyramine-modified silk fibroin hydrogels as a tunable platform for studying the interplay between progressive tissue mechanics and profibrotic biochemical signaling in vitro.

## Results and Discussion

### Tyramine modification of silk enables tunable, progressive hydrogel stiffening

Recapitulating the progressive mechanical stiffening of fibrotic tissue requires a chemically defined platform with controllable crosslinking kinetics. Silk fibroin (SF) was chemically modified with tyramine via EDC/NHS coupling chemistry to yield tyramine-modified silk fibroin (SF-TA, **Fig. 1A**)[33,36]. The tyramine modification adds additional phenol-containing crosslinking sites on top of the native tyrosine residues already present in the silk backbone. Successful tyramine conjugation to the SF backbone was confirmed by NMR spectroscopy (**Fig. S1**). The phenol groups serve as substrates for horseradish peroxidase (HRP) mediated oxidative crosslinking in the presence of hydrogen peroxide (H_2_O_2_), forming covalent di-tyramine bonds that produce a stable hydrogel network (**Fig. 1B**). Two formulations were prepared: pure SF (0%) and a 50:50 blend of SF and SF-TA (50%). Blending SF with SF-TA provided a straightforward means to control the extent of hydrogel crosslinking and enabled divergent compressive moduli profiles within a material class.

**Figure 1:**
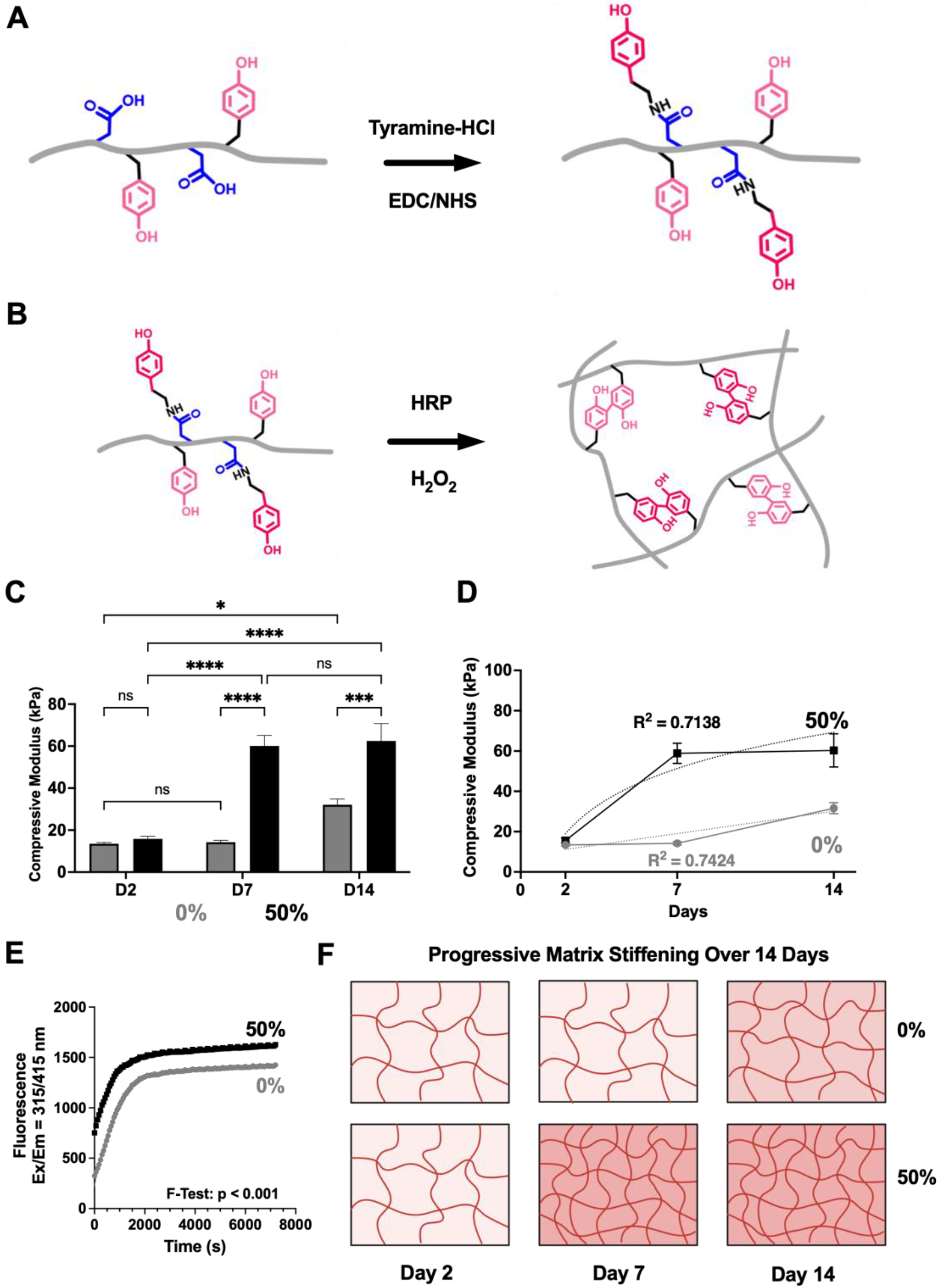
Tyramine modification of silk generates hydrogels with programmable mechanical stiffening, providing a tunable platform for fibrosis modeling. A) Silk is modified with tyramine-HCl via EDC/NHS chemistry to conjugate additional phenol crosslinking sites onto the silk backbone. B) Tyramine-modified silk is crosslinked via HRP/H_2_O_2_ enzymatic reaction to form a covalently crosslinked hydrogel. C) The compressive modulus of silk hydrogels cultured with NHLFs differs significantly between 0% (gray) and 50% (black) modified hydrogels across Day 7 and 14 timepoints. D) Stiffening trajectories over 14 days: the 0% SF-TA formulation stiffens linearly 0% (R^2^=0.7424), while the 50% SF-TA formulation follows a semi-logarithmic relationship (R^2^=0.7138). E) Gelation kinetics for 0% and 50% modified silk hydrogels (one-phase association model, F-test: p < 0.001). F) Schematic illustrating two models of cell-driven matrix stiffening: gradual (0%) and progressive (50%) over 14 days. N=5-6, Error bars = SEM. Two-way ANOVA with Tukey post-hoc test, * = p < 0.05, *** = p < 0.001, **** = p < 0.0001.

We asked whether these formulations produced distinct mechanical stiffening trajectories under a mono fibroblast cell culture condition. Normal Human Lung Fibroblasts (NHLFs) were seeded on top of 2 mm hydrogels, and compressive modulus was measured at days 2, 7, and 14 (**Fig. 1C**). Both formulations began at comparable stiffness at day 2 (0%: 13.5 kPa, 50%: 15.8 kPa), within the early fibrotic range [11–15]. The 50% formulation stiffened rapidly, reaching 60.1 kPa by day 7 and plateauing at 62.5 kPa by day 14, following a semi-logarithmic relationship over time (R^2^ = 0.71, **Fig. 1D**). The 0% SF-TA formulation stiffened gradually and continuously, reaching 14.3 kPa at day 7 and 32.1 kPa by day 14, following a linear relationship over time (R^2^ = 0.74, **Fig. 1D**). Prior characterization of TA hydrogels has demonstrated progressive stiffening over time, a behavior conserved in the presence of cells[33]. Gelation kinetics confirmed that the 50% SF-TA formulation reached a higher fluorescence plateau than the 0% SF-TA formulation, as fit by a one-phase association model (**Fig. 1E**, F-test p < 0.001). The rate constant K did not differ significantly between formulations (F-test, p = 0.5988), indicating that differences in crosslinking behavior reflect the extent of network formation rather than the reaction rate. These distinct stiffening profiles reflect the mechanical progression of early (0% SF-TA) and established (50% SF-TA) fibrotic lung tissue (**Fig. 1F**)[11–15], providing two temporally distinct mechanical environments for investigating fibroblast activation[5,9,16].

Matrix stiffness is a well-established driver of fibroblast activation and myofibroblast differentiation in fibrotic disease [5,9,16]. However, most in vitro models present static mechanical environments[3,15,23,24,37,38], failing to capture the progressive, time-dependent stiffening that characterizes fibrotic disease evolution in vivo[16,26–29]. By incorporating distinct stiffening trajectories, these two formulations introduce a temporal dimension to the mechanical modeling of fibrosis that is rarely investigated. This is particularly relevant given the concept of mechanical memory[30,31], whereby cells retain a functional imprint of prior mechanical environments that encodes lasting fibrogenic programs. Incorporating dynamic stiffening into fibrosis hydrogel models could more faithfully recapitulate the mechanical environment fibroblasts encounter during disease progression. This may further enable investigation into mechanical memory as a driver of persistent myofibroblast activation and position dynamic stiffening hydrogel platforms as tools for screening stage-specific antifibrotic therapeutics.

### Matrix stiffening attenuates NHLF metabolic activity without inducing senescence on 50% SF-TA hydrogels

Having established two mechanically distinct hydrogel platforms, we first sought to confirm their cytocompatibility for long-term fibroblast culture. These cells were cultured on 0% and 50% SF-TA hydrogels, with and without TGFβ (5 ng/mL), over 14 days, alongside tissue culture plate (TCP) controls (**Fig. 2A**). For each hydrogel treatment, 25,000 NHLF cells were added to 2 mm hydrogels in a 24-well transwell plate. These conditions remained constant throughout the study and were optimized to prevent cellular overgrowth while maintaining sufficient cell numbers for downstream analyses.

**Figure 2:**
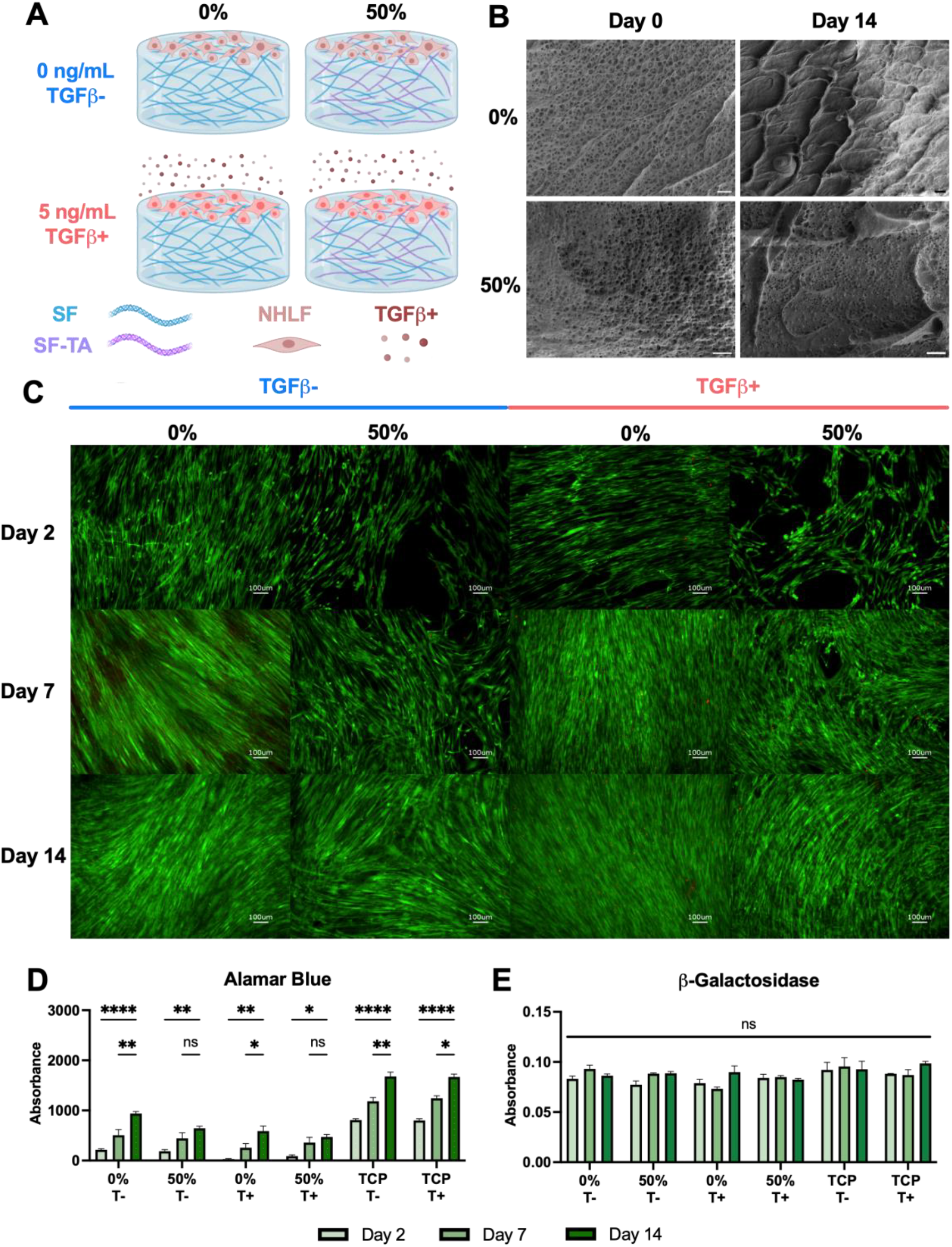
Tyramine-modified silk hydrogels are cytocompatible, yet cell proliferation plateaus without senescence maker elevation. A) Hydrogels were composed of either 0% or 50% mixture of tyramine-modified silk and were exposed to media without or with 5 ng/mL TGFβ. After hydrogels were cast, 25,000 NHLF cells were seeded on the surface. B) SEM imaging of the surface of the silk hydrogels incubated without cells for Day 0 and Day 14. C) Live/Dead imaging of Calcein AM for live cells (green) and Ethidium Homodimer-1 for dead cells (red) indicated cytocompatibility. D) Alamar Blue Assay for metabolic activity indicated that 0% gels showed an increase in activity over 14 days, while 50% gels plateaued from day 7 to 14. E) Mammalian β-Galactosidase Assay for senescence revealed no change for any timepoints tested over 14 days. N=3-4, Error bars = SEM.TCP = Tissue Culture Plate, T-= without TGFβ, T+ = 5 ng/mL of TGFβ. Two-way ANOVA, Tukey’s: ns = not significant p > 0.05, * = p < 0.05, ** = p < 0.01, *** = p < 0.001, **** = p < 0.0001.

To visualize the acellular hydrogel surface, SEM imaging revealed a porous surface architecture in both formulations at day 0, which closed over the 14-day culture period (**Fig. 2B**). Live/dead imaging confirmed high viability across all conditions and time points across three replicates (**Fig. 2C, Fig. S2-4**). TCP controls demonstrated similar viability across all time points and TGFβ conditions, confirming that neither culture duration nor TGFβ treatment induced high cytotoxicity (**Fig. S5-6**). The initial cell distribution observed on 50% SF-TA substrates was patchy, despite RGD functionalization; however, this resolved by day 7 and did not affect long-term culture outcomes. These data align with the literature, which indicates that silk is cytocompatible and requires cell adhesion assistance[36,39]. Qualitative evidence of cell elongation and spreading was observed over time, consistent across all conditions. The organized swirling cell distribution patterns observed qualitatively across substrates resemble those reported in Normal Human Bronchial Epithelial (NHBE)/NHLF co-cultures by *Barron et al*.[38].

Late-stage fibrosis is characterized by myofibroblast persistence, which is associated with reduced proliferative metabolic activity as matrix-consolidating cells exit the cell cycle and resist apoptotic resolution[5,19]. Metabolic activity was assessed using Alamar Blue and found to plateau between day 7 and day 14 in the 50% SF-TA, both with and without TGFβ. In contrast, there was a continuous increase in 0% SF-TA substrates and TCP, regardless of TGFβ treatment (**Fig. 2D**). TCP controls reached approximately 2-fold higher metabolic activity than hydrogel conditions by day 14, consistent with other literature reports[40,41]. To determine whether the plateau reflected cellular senescence, β-galactosidase activity was quantified across all conditions and time points (**Fig. 2E**). Values showed no significant differences. The absence of elevated β-galactosidase activity suggests that the metabolic plateau observed with 50% SF-TA substrates reflects a persistent myofibroblast state.

Together, these findings suggest that 0% and 50% SF-TA hydrogels support mechanically distinct fibroblast states consistent with early and late fibrosis, respectively. Alongside the cytocompatibility data, the continuous metabolic expansion observed on softer substrates reflects a proliferative, active fibroblast phenotype, while the plateau observed on stiffer substrates is consistent with the myofibroblast persistence characteristic of advanced disease[2,5]. Capturing both states within a single biomaterial system provides a foundation for interrogating stage-dependent fibroblast behavior. This is an important consideration given that anti-fibrotic interventions are known to elicit distinct responses depending on disease stage [2,5,42].

### NHLFs cultured on 0% SF-TA hydrogels display an inflammatory cytokine profile consistent with early fibrotic signaling

Fibroblast secretion of inflammatory cytokines is a hallmark of early fibrotic disease, reflecting the onset of the wound-healing cycle[2,5,43]. In fibrosis, this cycle fails to resolve, perpetuating dysregulated inflammatory signaling [2,5,43]. Fibroblasts secrete several cytokines, IL-6[43], IL-8[44,45], and MCP-1[46,47], which drive macrophage recruitment and perpetuate a pro-inflammatory microenvironment[2,5]. This pro-inflammatory environment promotes collagen deposition via two fibrosis-prominent pathways: TGFβ and Wnt[1,48,49]. In parallel, matrix stiffness can independently drive collagen production via mechanotransduction [2,5,29]. We asked whether progressive matrix stiffening drives differential inflammatory signaling and if that immune signaling impacts collagen secretion.

To investigate the immune profiles of our fibroblast cultures, a 13-plex LEGENDplex cytokine panel was used to profile across all conditions and timepoints (**Fig. 3A**). Media controls confirmed that cytokine signal was entirely cell-derived (**Fig. S7A-C**). Of the 13 analytes measured, only IL-6, IL-8, and MCP-1 produced detectable signals across experimental conditions (**Fig. 3A**). The levels of TNF-α, IFN-γ, and IL-1β remained at or near the limit of detection across all conditions (**Fig. 3A**). A recent study with IPF human fibroblasts obtained from explants showed elevated levels of TNF-α and IFN-γ[50]. However, this study had 10 times as many cells, and other studies report levels similar to ours in cell culture[38]. Fibroblast secretion of IL-1β is typically associated with innate immune responses to foreign material or pathogen recognition, rather than fibroblast-intrinsic signaling[51–54]. The remaining six analytes were indistinguishable from controls across all conditions and timepoints. This was expected, as these cytokines are predominantly secreted by immune cells rather than fibroblasts[2,43,47,55,56].

**Figure 3:**
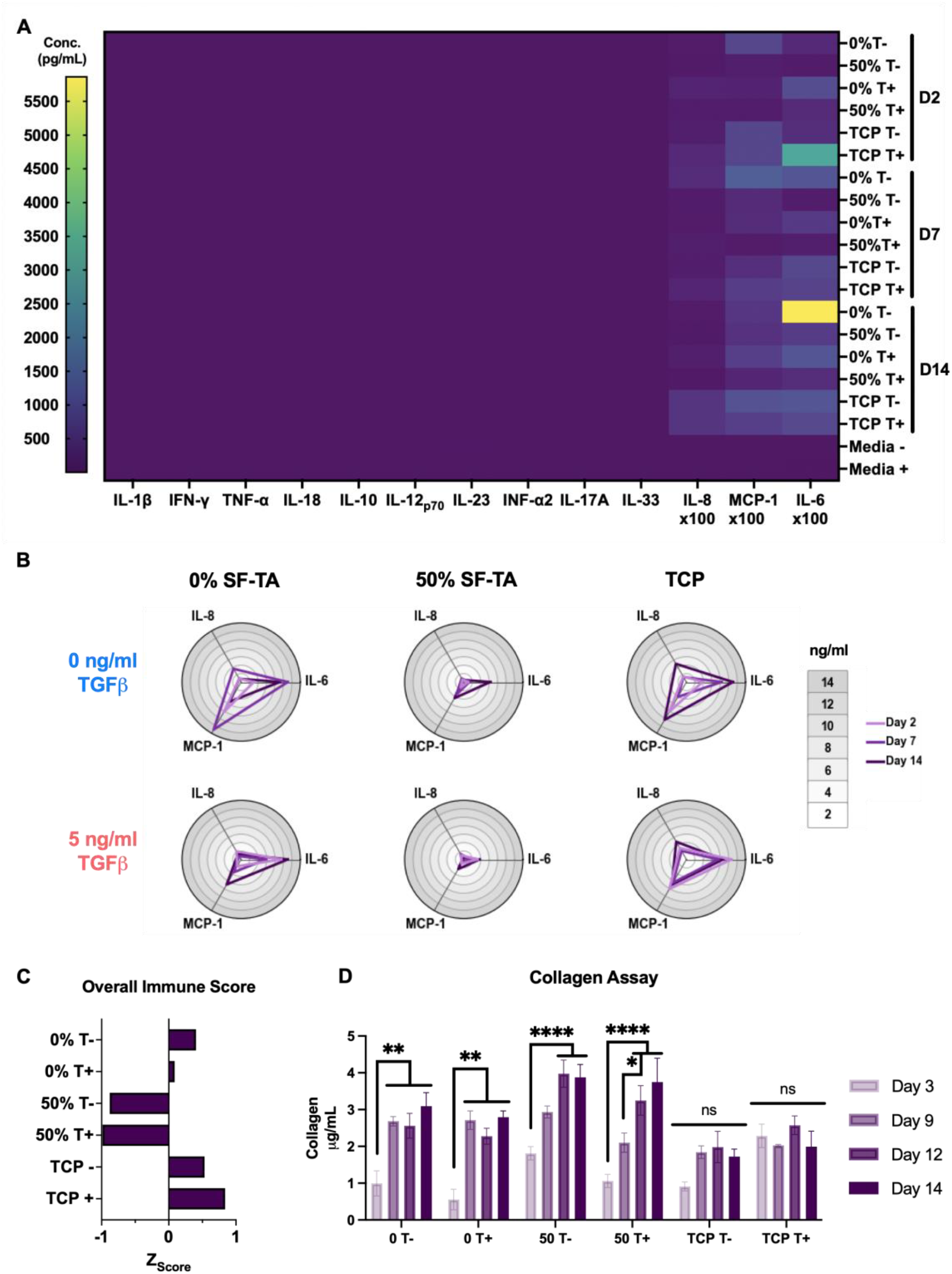
Dynamic stiffening alters immune profiles, with 0% hydrogels fostering an immunostimulatory environment. Cell culture media were obtained on Day 2, Day 7, and Day 14, and a human Legendplex panel was conducted. A) Heatmap of cytokine concentrations (pg/mL) across all conditions and timepoints, revealing that IL-6, MCP-1, and IL-8 are the consistently detectable analytes. B) Radar plots of IL-6, MCP-1, and IL-8 levels across timepoints with 0% SF-TA and TCP showing higher cytokine levels compared to 50% SF-TA. C) Overall immune Z-score indicates that 0% SF-TA and TCP conditions are immunostimulatory relative to 50% SF-TA hydrogels. D) Secreted collagen content (µg/mL) from 0% SF-TA, 50% SF-TA, and TCP conditions with and without TGFβ at Day 3, Day 9, Day 12, and Day 14. N=3-4, Error bars = SEM. TCP = Tissue Culture Plate, T-= without TGFβ, T+ = 5 ng/mL TGFβ. Two-way ANOVA, Tukey’s: ns = not significant p > 0.05, * = p < 0.05, ** = p < 0.01, **** = p < 0.0001.

Breaking down the inflammatory signal further, 0% SF-TA substrates produced higher overall cytokine secretion than 50% SF-TA substrates (**Fig. 3B**). The 0% SF-TA substrates were also similar to TCP controls. To compare relative inflammatory response across conditions, Z-score analysis was used to normalize cytokine levels to the mean across all conditions. Overall (**Fig. 3C**) and individual cytokine (**Fig. S7D-F**) immune scores were calculated for IL-6, IL-8, and MCP-1 across the three time points. Together, the Z-score shows that TCP conditions ranked highest, followed by 0% and 50% SF-TA. Of note, the 50% SF-TA always had a negative Z-score. The literature supports that stiffer substrates, excluding excessively stiff TCP, can suppress inflammatory signaling[56,57]. The overall Z-scores were more strongly separated by substrate condition than by TGFβ treatment, indicating that the substrate and its stiffness were the dominant drivers of the inflammatory response.

Divergent cytokine signaling can drive differential collagen deposition[2,5,43,58]. Collagen secretion was therefore assessed to determine whether the inflammatory profiles were reflected in matrix output. The TCP control conditions showed no significant change in collagen over the 14 days (**Fig. 3D**). On 0% SF-TA substrates, collagen levels increased significantly from day 3 to day 9 and plateaued. In contrast, the 50% SF-TA substrates showed a gradual increase in collagen production, reaching steady levels by day 12. Notably, TGFβ treatment did not alter collagen output in either condition, suggesting that neither trajectory was primarily TGFβ-dependent. This could be an effect of the 5 ng/mL concentration used or of the fact that TGFβ was added as an exogenous media supplement[17,18,26]. However, the different kinetic profiles of collagen secretion do suggest a divergence in regulatory mechanisms. The early burst and sustained collagen secretion observed on 0% SF-TA substrates may reflect the immunostimulatory environment, as the cytokines measured are implicated in Wnt-pathway-driven collagen deposition[59,60]. The 50% SF-TA substrates lacked the same immune-stimulatory profile; however, collagen levels increased, indicative of mechanotransduction-driven collagen production [2,5,29]. Despite these mechanistically distinct trajectories, day 14 secreted collagen levels were equivalent across the 0% and 50% SF-TA substrates. This implies comparable secretory capacity across divergent mechanical and inflammatory microenvironments. These results suggest that matrix stiffness can drive collagen secretion to levels comparable to those achieved through cytokine-mediated signaling.

The data on metabolic, cytokine, and collagen together advance the proposal that 0% and 50% SF-TA substrates drive mechanistically distinct fibrogenic phenotypes. On 0% SF-TA substrates, continuous metabolic expansion, elevated pro-inflammatory cytokine secretion, and quick collagen secretion reflect an early fibrotic state[1,2]. For the 50% SF-TA substrates, there is plateaued metabolic activity and attenuated inflammatory cytokine output, while having comparable collagen accumulation, more consistent with a late fibrotic state[15,16,21].

### Progressive matrix stiffening and TGFβ synergistically drive α-SMA stress fiber incorporation in 50% SF-TA hydrogels

The substrate-dependent differences in inflammatory signaling and collagen deposition raised the question of whether fibroblast cytoskeletal organization was similarly affected. Myofibroblast differentiation is defined by the incorporation of α-SMA into contractile stress fibers[5,16,19,61]. Cytoskeletal organization was assessed qualitatively by immunofluorescence staining for F-actin and α-SMA across all four conditions at days 2, 7, and 14, with consistent imaging thresholds applied across days (**Fig. 4, Fig. S8**). Individual channel images for each time point are provided in the supplement (**Fig. S9-11**).

**Figure 4:**
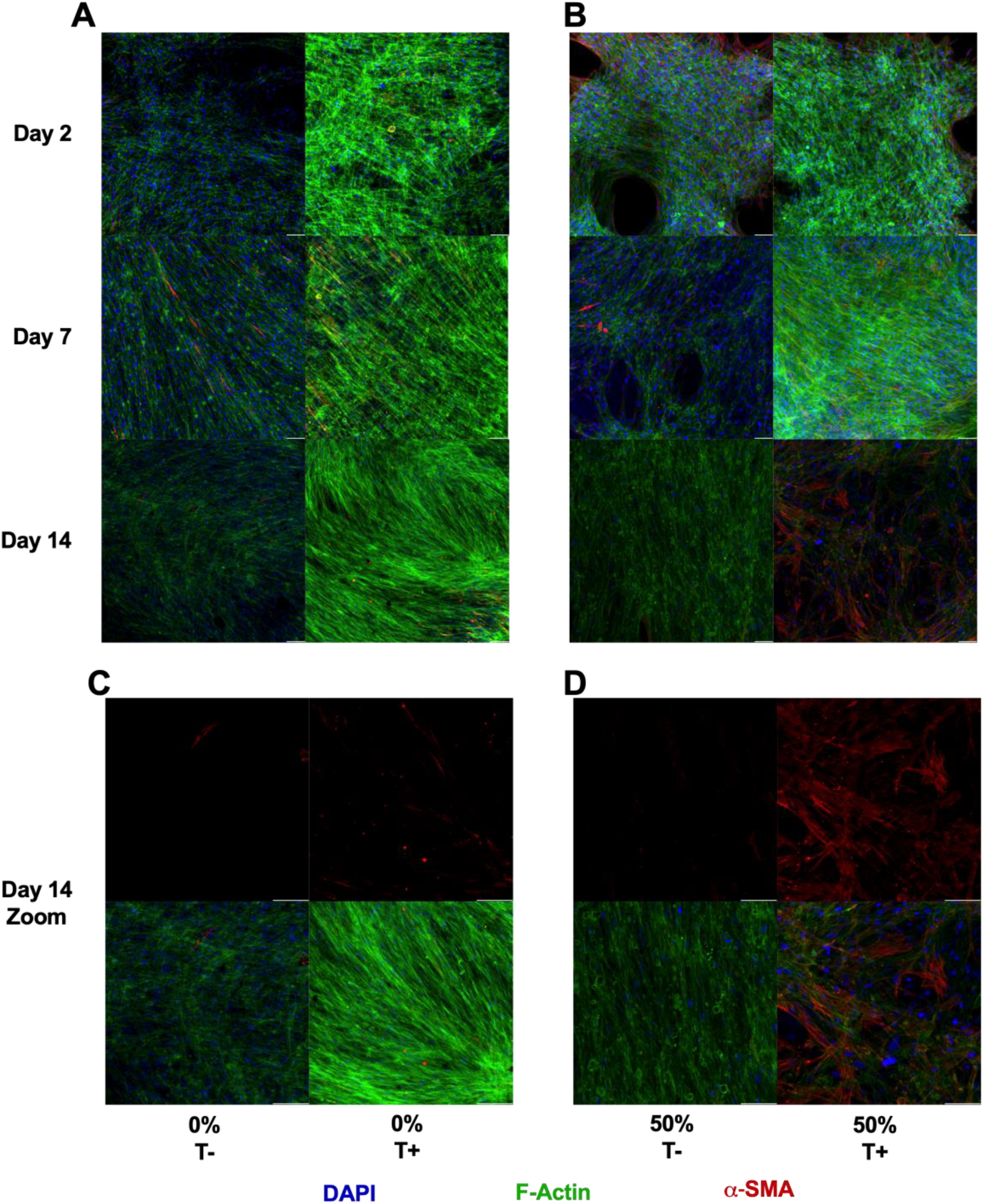
Matrix stiffness and TGFβ drive myofibroblast activation as visualized by immunofluorescence. Immunofluorescence images of NHLFs stained for DAPI (blue), F-Actin (green), and α-SMA (red) were obtained for Day 2, Day 7, and Day 14 for treatments with and without TGFβ. A) 0% SF-TA, B) 50% SF-TA. C) Zoomed in Day 14 images of 0% SF-TA conditions. D) Zoomed in Day 14 images of 50% SF-TA conditions. T-= without TGFβ, T+ = 5 ng/mL TGFβ. Scale bar = 100 µm.

Progressive cytoskeletal reorganization was observed across all conditions over the 14-day culture period. On 0% SF-TA substrates, cells displayed predominantly F-actin cytoskeletal organization with qualitatively scarce α-SMA signal at day 2 (**Fig. 4A**). Sparse stress fibers emerged by day 7, and more prominently in the TGFβ-treated condition. The minimal α-SMA signal remained at day 14 regardless of TGFβ treatment (**Fig. 4A**). On 50% SF-TA substrates, qualitative α-SMA signal was present at day 2 even in the absence of TGFβ, suggesting that matrix stiffness alone may initiate early α-SMA expression (**Fig. 4B**). By day 14, the TGFβ-treated condition displayed visibly denser and more organized α-SMA-positive stress fibers co-localizing with F-actin (**Fig. 4B-D**). These observations support that synergistic mechanical and TGFβ signaling promote the cytoskeletal organization of persistent myofibroblast-associated phenotypes [3,16,19,21].

These findings further support the distinction between early and late fibrotic states modeled by 0% and 50% SF-TA substrates, respectively. Despite elevated inflammatory cytokine secretion and a gradual linear stiffening profile on 0% SF-TA substrates, this microenvironment was insufficient to drive consistent α-SMA stress fiber incorporation, highlighting the requirement for mechanotransductive feedback in myofibroblast commitment[16,61]. Within the 50% SF-TA condition, matrix stiffness alone initiated early α-SMA expression, but sustained and organized stress fiber incorporation required TGFβ co-stimulation, suggesting that both mechanical and biochemical stimuli may be necessary to recapitulate late-stage fibrotic disease in vitro[16,21].

### Synergistic mechanical and biochemical signaling drives myofibroblast gene expression in 50% SF-TA hydrogels treated with TGFβ

Myofibroblast differentiation is governed by both mechanical and biochemical cues, with TGFβ and matrix stiffness acting as key drivers of fibroblast activation (**Fig. 5A**)[16–18,21]. Matrix stiffness is sensed through integrin receptors, particularly Integrin Beta-1 (ITGB1), which clusters at focal adhesion complex[5]. One prominent component of this complex is Focal Adhesion Kinase (FAK)[2,62]. Several pathways can be activated through focal adhesion complex signaling: Ras Homolog Family Member A (RhoA), Rho-associated protein kinase (ROCK), and Yes-Associated Protein (YAP)[29,63]. Of note, RhoA is ROCK-associated; however, RhoA can also be initiated independently through cell-surface Receptor Tyrosine Kinase (RTK) pathways[29,62,64,65]. YAP forms complexes and promotes the production of Alpha-Smooth Muscle Actin (ACTA2) and Connective Tissue Growth Factor (CTGF)[29,63,66]. Meanwhile, ROCK promotes nuclear translocation Myocardin-Related Transcription Factor-A (MRTF-A), which also drives ACTA2 and CTGF production[5,62,65]. Importantly, the RTK pathway forms a positive feedback loop with CTGF[65]. In parallel, TGFβ signals through Smad-dependent pathways to drive expression of ACTA2, CTGF, and Collagen Type I Alpha 1 (COL1A1)[17,18,26,59]. Both RTK and ITGB1 signaling can also drive collagen synthesis[2,5,67]. The transcriptional response to progressive stiffening and TGFβ treatment was evaluated across a panel of nine genes targeting key nodes of these pathways by TaqMan qPCR, normalized to day 2 within each condition (**Fig. 5B**).

**Figure 5:**
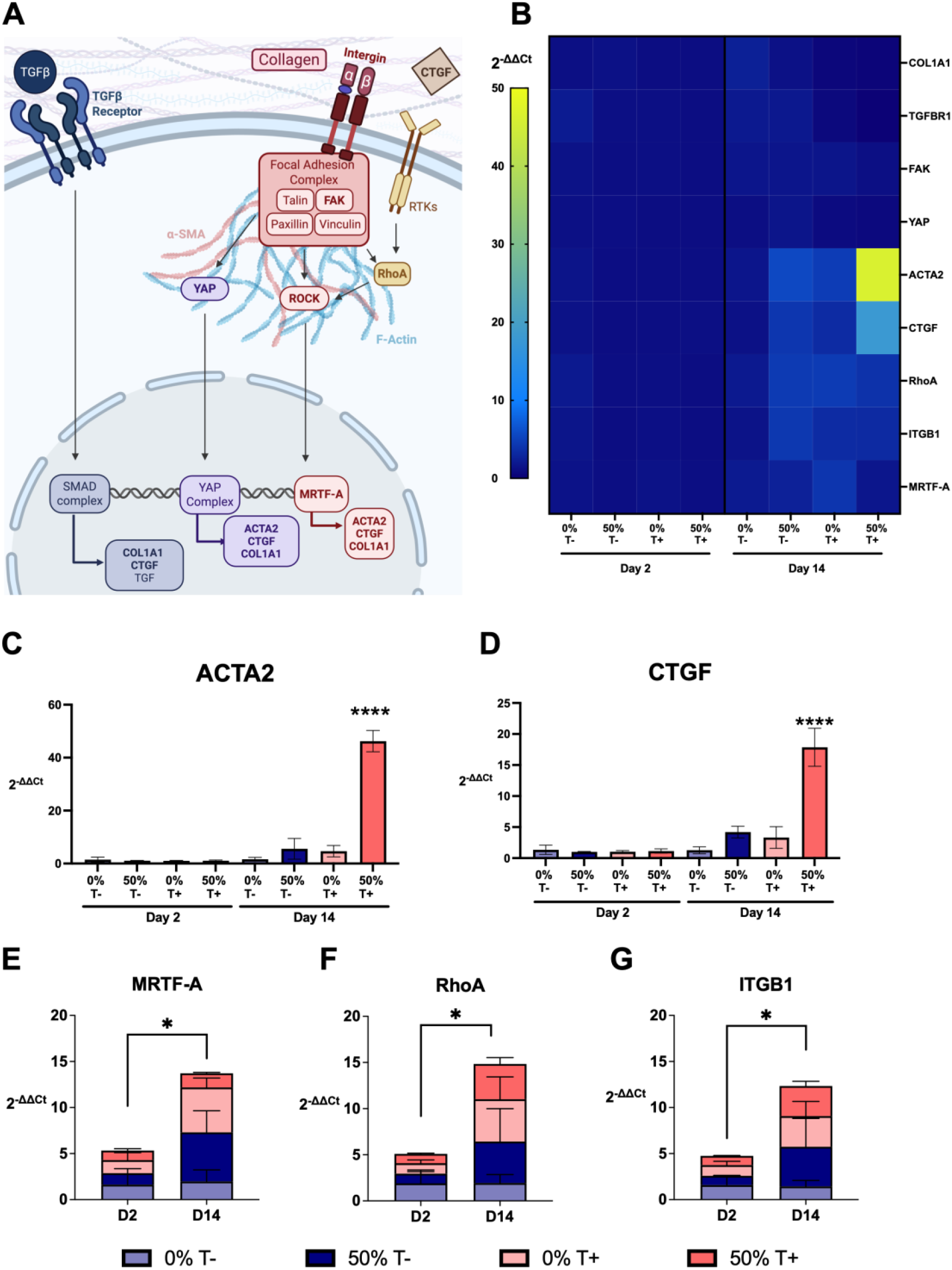
Gene expression profiling of mechanosensing and fibrotic pathways demonstrates that rapid stiffening synergizes with TGFβ stimulation to upregulate fibrotic markers at Day 14. A) Schematic of the mechanosensitive and profibrotic signaling pathways targeted by gene expression analysis. B) Heatmap of RT-qPCR gene expression, normalized to GAPDH and to the Day 2 average for each group, reported as 2^-ΔΔCt^ for mechanosensitive and profibrotic genes. C) ACTA2 (α-smooth muscle actin) expression is significantly elevated in 50% with TGFβ conditions at Day 14. D) CTGF (connective tissue growth factor) expression mirrors this response, with significant upregulation in 50% with TGFβ hydrogels at Day 14. Global trends in E) MRTF-A, F) RhoA, and G) ITGB1 reveal significant upregulation from Day 2 to Day 14, suggesting engagement of mechanosensitive and profibrotic pathways across all conditions. T-= without TGFβ, T+ = 5 ng/mL TGFβ. N=3, Error bars = SEM. Two-way ANOVA, Tukey’s (C-D), Welch’s t-test (E-G), ns = not significant, p > 0.05, * = p < 0.05, **** = p < 0.0001.

At day 14, significant upregulation of both ACTA2 and CTGF was observed in the 50% SF-TA condition treated with TGFβ (p<0.0001, **Fig. 5C-D**). Neither mechanical stiffening alone nor TGFβ treatment on soft substrates was sufficient to drive significant upregulation of either gene. This needed synergy is consistent with the known cooperation between Smad-dependent TGFβ signaling and mechanosensitive transcriptional programs [5,18,26]. Existing fibrosis models often rely on multi-factor biochemical cocktails [68] or materials derived from decellularized extracellular matrix[3] to drive myofibroblast gene expression. In a 2D static stiffness model, fibrotic-range stiffness alone was sufficient to upregulate ACTA2, but CTGF remained unchanged[15]. The in vitro system presented in this paper advances fibrotic tissue modeling by incorporating progressive mechanical stiffening with TGFβ, which, together, were sufficient to drive upregulation of both ACTA2 and CTGF. More research is needed to fully elucidate whether dynamic stiffening history, rather than endpoint modulus alone, is a critical determinant of full myofibroblast transcriptional commitment [30,31]. However, growing evidence implicates mechanical memory as a driver of persistent fibrogenic programming, suggesting its consequence[69–71]. The progressive stiffening platform presented here provides a tool for further interrogation of these questions. Furthermore, the chemically defined and tunable nature of this platform provides a simplified foundation for introducing multicellular complexity. Thus, enabling future investigation of crosstalk between fibroblasts, immune, or epithelial cell populations.

While ACTA2 and CTGF showed condition-dependent upregulation, COL1A1 transcript levels showed no significant changes between day 2 and day 14 across any condition, or when pooled (**Fig. 5B, Fig. S10**). This lack of gene expression change contrasted with the continued collagen protein secretion observed across substrate conditions. This disconnect may reflect post-transcriptional regulation of collagen synthesis, reduced activity of matrix-degrading pathways, or contributions from additional collagen subtypes not captured by COL1A1 alone [2,5,62,67]. We hypothesize that the total collagen measured reflects contributions from other collagen isoforms, which are known to change across the fibrosis timeline [2,5,67]. Resolving the relationship between collagen gene expression and protein deposition represents an opportunity for future mechanistic investigation using this platform.

When analyzed individually, ITGB1, RhoA, and MRTF-A showed no significant changes between day 2 and day 14 within any single condition (**Fig. 5B**). However, when pooled across all conditions, all three genes increased significantly from day 2 to day 14 (Welch’s t-test, p<0.05; **Fig. 5E-G**), suggesting a consistent, but modest, time-dependent upregulation of ITGB1/RhoA/MRTF-A. This global trend is consistent with a mechanical priming effect in which fibroblasts progressively engage cytoskeletal tension signaling over the culture period[2,5,71]. In contrast, FAK and YAP transcript levels remained flat across all conditions and timepoints (**Fig. S10**). Both MRTF-A and YAP serve as mechanosensitive transcriptional regulators; however, they have divergent global trends. This could be due to MRTF-A being more directly associated with stiffness-induced myofibroblast differentiation and ACTA2 expression[2,19,65], while YAP is more broadly implicated in fibroblast proliferation and ECM synthesis[66,72]. FAK is also an immediate-early responder to integrin engagement and matrix stiffness, becoming active upon initial cell seeding and substrate contact[73,74]. As samples were normalized to day 2 within each condition, any stiffness-dependent differences in FAK activation that occurred prior to or at the earliest time point would not be captured by this analysis. The global upregulation of ITGB1 suggests that extended culture periods or earlier collection timepoints may be necessary to fully capture the temporal dynamics of FAK activation and its downstream consequences for mechanotransductive signaling in this system.

In fibrosis, the role of TGFβ and its receptor, TGFB1, is well documented[17,18,26]. Interestingly, TGFB1 transcript levels did not differ significantly between day 2 and day 14 across any condition or when pooled (**Fig. 5B, Fig. S10**). This is notable given that TGFβ treatment was clearly required to drive significant upregulation of ACTA2 and CTGF and to commit α-SMA stress fiber formation in the imaging data. The absence of TGFB1 transcriptional change may reflect that the 5 ng/mL exogenous dose used in this study was insufficient to drive detectable changes in expression levels, or that dynamic TGFβ concentrations may be required to elicit transcriptional responses. In vivo, TGFβ is expressed by immune cells, particularly macrophages, which are absent in this monoculture system[17,18,59]. Future multicellular models incorporating macrophages or broader immune cell stimulation may therefore reveal more dynamic TGFB1 regulation and provide insight into the interplay between inflammatory and mechanotransductive signaling in fibrosis progression.

These gene expression data provide the final layer of evidence for the two-state fibrosis model. Global upregulation of ITGB1, RhoA, and MRTF-A across all conditions reflects time-dependent mechanical priming independent of substrate stiffness or TGFβ treatment. While exclusive upregulation of ACTA2 and CTGF in 50% SF-TA substrates treated with TGFβ demonstrates that myofibroblast transcriptional commitment requires the convergence of both mechanical and biochemical signals. This platform, capturing both the inflammatory and mechanotransductive axes of fibrotic progression, provides a foundation for future investigation of stage-specific therapeutic targets and multicellular interactions across the fibrotic disease continuum.

### SF-TA hydrogels recapitulate early and established fibrosis states in vitro

Fibrosis progresses through distinct stages, from early inflammatory fibroblast activation to established myofibroblast persistence. The data presented here support the case that 0% and 50% SF-TA hydrogels are consistent with features of these two stages, respectively (**Fig. 6**). On 0% SF-TA substrates, fibroblasts remained proliferative and metabolically active, secreting IL-6, IL-8, and MCP-1 consistent with early inflammatory fibroblast activation and macrophage recruitment [2,5,43]. Alongside this, collagen secretion is observed, consistent with cytokine-mediated matrix deposition [1,62,71]. On 50% SF-TA substrates, fibroblasts exhibited a metabolic plateau without senescence and suppressed inflammatory cytokine secretion [5,19]. In the presence of TGFβ, ACTA2 and CTGF were significantly upregulated, along with α-SMA stress fiber incorporation, indicative of myofibroblast activation and persistence [5,19]. Furthermore, collagen secretion continued throughout culture [62,67]. Mechanical stiffness appears to drive fibroblast fate, with TGFβ serving as a necessary co-stimulus for full myofibroblast commitment.

**Figure 6.**
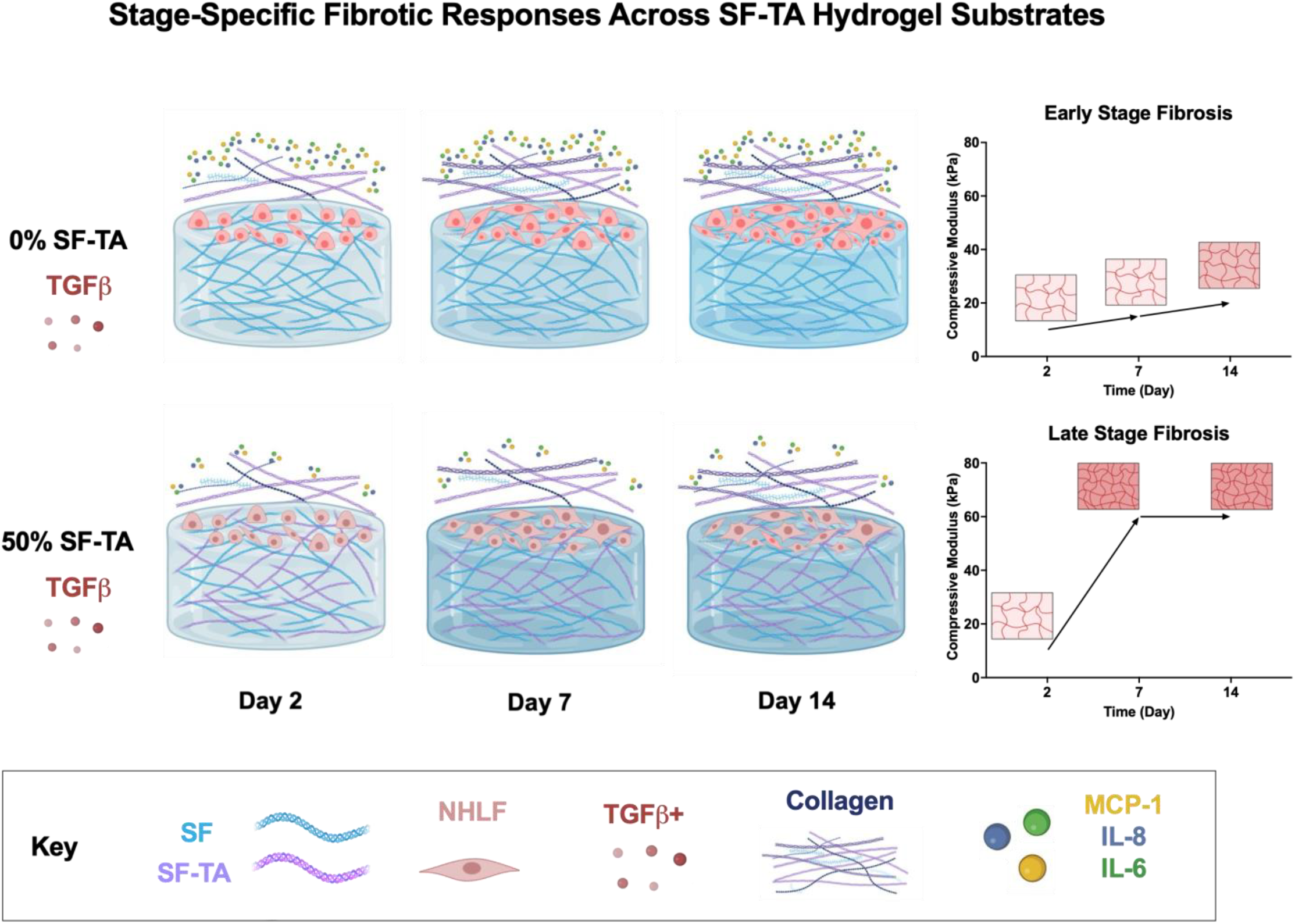
Schematic summary of stage-specific fibrotic responses across 0% and 50% SF-TA hydrogel substrates. Top row: NHLFs on 0% SF-TA hydrogels at Day 2, Day 7, and Day 14, exhibiting a proliferative, inflammatory phenotype consistent with early fibroblast activation, characterized by secretion of IL-6, IL-8, and MCP-1 (dots) and progressive collagen deposition (fibers). Bottom row: NHLFs on 50% SF-TA hydrogels at Day 2, Day 7, and Day 14, exhibiting metabolic plateau and lower inflammatory cytokine secretion, gradual collagen secretion, with TGFβ-dependent myofibroblast activation. Together, these two substrate conditions recapitulate distinct stages of the fibrotic disease continuum.

As with all in vitro models, limitations exist; these silk hydrogels represent a simplification of the fibrotic microenvironment, and several directions remain open for further development. Fibroblasts are currently cultured on the gel surface rather than within it. Transitioning toward 3D encapsulation would more closely replicate the mechanical environment fibroblasts experience in vivo. Several studies have demonstrated that this distinction meaningfully affects downstream outcomes, including myofibroblast differentiation[3]. Achieving this with SF-TA would require optimizing gelation kinetics and compressive moduli to enable adequate cell spreading. The use of exogenous TGFβ similarly points toward future co-culture models incorporating macrophages, epithelial cells, or other stromal populations to better capture endogenous cytokine dynamics. The tunability of the SF-TA stiffening profiles provides a foundation for future investigations into mechanical memory, multicellular systems, and their impact on fibrogenic programs. Together, these findings position SF-TA hydrogels as a tunable mechanobiology platform for investigating stage-specific fibrotic mechanisms and screening therapeutic strategies across the fibrotic disease continuum.

## Conclusions

In summary, tyramine-modified silk fibroin hydrogels with tunable stiffness provide a cytocompatible platform for in vitro modeling of distinct stages of pulmonary fibrosis. By exploiting the progressive stiffening profiles of 0% and 50% SF-TA formulations, this system captures the early inflammatory fibroblast activation and established myofibroblast persistence state within a single biomaterial platform. The convergence of metabolic, cytokine, collagen, cytoskeletal, and gene expression data across both conditions provides multi-parametric support for the distinct stage-associated fibrogenic phenotypes and demonstrates the platform’s sensitivity to substrate- and treatment-dependent fibroblast behavior. The synergistic interaction between progressive mechanical stiffening and TGFβ in driving myofibroblast commitment highlights the importance of incorporating both mechanical and biochemical cues in fibrosis modeling. The distinct stiffening trajectories of 0% and 50% SF-TA hydrogels further provide a tunable foundation for investigating mechanical memory and how prior mechanical history encodes lasting fibrogenic programs. Future work incorporating multicellular complexity, including immune cell populations, will further extend the utility of this platform for investigating the fibrotic disease continuum and evaluating stage-specific therapeutic strategies.

## Methods

### Preparation of Silk Fibroin

Aqueous silk fibroin (hereafter termed silk) solutions were prepared as previously described [36]. In short, *Bombyx mori* cocoons (Tajima Shoji Co., Ltd., Yokohama, Japan) were degummed to remove sericin proteins by boiling 5 g of cut cocoons in 2 L of 0.02 M sodium carbonate solution (Sigma-Aldrich, Burlington, MA, USA) for 60 mins, followed by thorough rinsing in deionized water. The degummed fibers were allowed to dry overnight, then solubilized in a 25% (w/v) lithium bromide (Sigma-Aldrich) solution at 9.3 M for 4 hrs at 60°C. The solution was then dialyzed against distilled water using regenerated cellulose dialysis tubing (MWCO: 3,500 Da, Spectra/Por® 3 Standard RC Tubing, Spectrum Laboratories Inc., Rancho Dominguez, CA, USA). Over 3 days, the dialysis water was changed 6 times. This solution was then centrifuged to remove insoluble particulates, and the concentration was calculated by measuring the mass of silk remaining after drying a known volume of the aqueous solution.

### Synthesis of Silk Fibroin-Tyramine (SF-TA)

SF-TA was synthesized via carbodiimide-mediated reactions as previously described [33,36]. Briefly, 2% (w/v) Silk was prepared in 0.05 M 2-(N-morpholino) ethanesulfonic acid (MES, Thermo Scientific, Waltham, MA, USA) buffer (pH 6.0) and mixed with tyramine hydrochloride (Tyr-HCl, Sigma-Aldrich) at a ratio of 500 mg Tyr-HCl: 1 g Silk protein. The reaction was initiated by the addition of 1-ethyl-3-(3-dimethalylaminopropyl) carbodiimide hydrochloride (EDC, Sigma-Aldrich) and N-hydroxysuccinimide (NHS, Sigma-Aldrich). The solution was gently stirred for 18 hrs. at room temperature, then filtered through a 40 μm cell strainer and dialyzed against DI water using a 3.5 kDa MWCO for 3 days with 6 water changes. Functionalization of silk-tyramine was characterized by Nuclear magnetic resonance (NMR) by reconstituting in deuterated water at a concentration of 10-15 mg mL^-1^. NMR was evaluated using Topspin software.

### Hydrogel fabrication

Silk hydrogels were prepared to a final total polymer concentration of 4% (w/v) using mixtures of nonfunctionalized silk fibroin (NSF) and tyramine-modified silk fibroin (SF-TA). Two formulations were generated: 0% SF-TA hydrogels containing 40 mg/mL NSF and 0 mg/mL SF-TA, and 50% SF-TA hydrogels containing 20 mg/mL NSF and 20 mg/mL SF-TA. All hydrogels were formulated with final concentrations of 0.005% H_2_O_2_ (Sigma-Aldrich), 10 U/mL Horseradish Peroxidase (HRP, Type-1, Sigma-Aldrich), 1x HD Buffer, and 0.1 mM Cyclo (-RGDyK) (RGD, AnaSpec, Fremont, CA, USA), with Invitrogen™ UltraPure™ DNase/RNase-Free Distilled Water (Thermo Scientific) added to the remaining volume. The 1x HD Buffer was made with 1 M Corning™ HEPES (Corning Life Sciences, Tewksbury, MA, USA) and D-(+)-Glucose (Sigma-Aldrich), with a final concentration of 40 mM HEPES and 5% D-(+)-Glucose. For sterile conditions, silk was filtered using a Millipore Steriflip Vacuum Tube Top Filter, pore size 0.22 μm (Sigma-Aldrich). The hydrogel precursor solution (NSF, SF-TA, HD Buffer, RGD, water) was made, and then HRP and H_2_O_2_ were added last. Hydrogels were then cast.

### Gelation kinetics

Hydrogel precursor solutions were mixed immediately before analysis, and 200 μL was aliquoted into a 96-well plate for kinetic measurement. To initiate gelation, 1 μL H_2_O_2_ was added, and the mixture was pipet-mixed right before reading. Gelation was monitored for 2 hrs. using a plate reader by tracking dityrosine fluorescence generated during enzymatic crosslinking. Fluorescence was measured at Ex/Em = 320/410 nm at 37 °C in kinetic mode over the course of gel formation (Thermo Scientific Varioskan Lux). Data were analyzed with GraphPad Prism (Boston, MA, USA) using a one-phase association model and an extra-sum-of-squares F Test with p < 0.05.

### Cell culture

Normal human lung fibroblasts (NHLFs, PCS-201-013, ATCC, Manassas, VA, USA) were cultured in Dulbecco’s Modified Eagle Medium, high glucose (DMEM, ThermoFisher) supplemented with 10% Gibco Fetal Bovine Serum, Premium (FBS, ThermoFisher), 1% Gibco non-essential amino acids (ThermoFisher), and 1% Gibco antibiotic–antimycotic (ThermoFisher). Cells were maintained in a humidified incubator at 37°C with 5% CO_2_, and the culture medium was replaced every 2-3 days. Culture media was supplemented with fibroblast growth Gibco factor-2 (FGF-2, ThermoFisher) by adding 5 pg/mL. For cell passaging, Gibco 0.05% trypsin-EDTA (ThermoFisher) and Gibco DPBS (ThermoFisher) were used.

### Hydrogel cell seeding

For cell-based experiments, hydrogels were cast into Falcon 24-well plate transwell inserts with a 0.4 µm transparent PET membrane (Corning Life Sciences) to a thickness of 2 mm, and allowed to gel for 1 h at 37°C and 5% CO_2_. Gels were then incubated in culture media for an additional 2 h prior to cell seeding. Cells were seeded onto each hydrogel at a density of 25,000 cells per well. A total of 1 mL of media was given to each well: 300 μL apical, 700 μL basolateral. For treated conditions, transforming growth factor-β (rhTGF-β1, R&D Systems, Minneapolis, MN, USA) was added to the culture media at a final concentration of 5 ng/mL.

### Unconfined Mechanical Compression

Unconfined compression was performed on a TA Instruments Discovery HR20 rheometer (TA Instruments, New Castle, DE, USA) between stainless steel parallel plates. Hydrogels were excised from the transwell using a 6 mm biopsy punch. The hydrogels were placed under a preload of ∼0.05 N to ensure complete surface contact. Two load-unload cycles to 15% strain at a rate of 0.5% per second, and the second cycle was used for analysis to eliminate artifacts. All moduli were calculated from the slope of the stress-strain curves using up to 5% strain. Hydrogels were cast as described above, and cells were seeded and maintained at 37°C and 5% CO_2_, with media changed every 2-3 days until the timepoint evaluation. Data were analyzed using custom Python code, cross-validated against Microsoft Excel (Redmond, WA, USA) outputs, and graphed in GraphPad Prism.

### Scanning electron microscopy (SEM)

Hydrogels were incubated in culture media for 3 h to allow media absorption prior to structural analysis. Samples were then flash frozen by immersion in liquid nitrogen. Frozen samples were transferred to -80°C and subsequently lyophilized (Labconco, Kansas City, MO, USA) for 24 h. To obtain cross-sections, lyophilized hydrogels were immersed in liquid nitrogen for ∼2 min to increase brittleness and then fractured. Samples were mounted onto aluminum SEM stubs using conductive carbon tape and sputter-coated with platinum prior to imaging. Microstructures were imaged using a Zeiss EVO MA10 Scanning Electron Microscope (Oberkochen, Germany) at an accelerating voltage of 5 kV. Scale bars were added in ImageJ (National Institutes of Health (NIH), Bethesda, MD, USA).

### Live/Dead viability staining

Cell viability was assessed using a Live/Dead staining assay with calcein AM (ThermoFisher) and ethidium homodimer-1 (ThermoFisher). Samples were washed and then incubated with staining solution prepared in DPBS containing calcein AM (Live, 1:500) and ethidium homodimer-1 (Dead, 1:200) for 20 min at 37°C and 5% CO_2_. After staining, hydrogels were carefully punched out of the culture wells, flipped cell-surface faced down, and placed onto 6-well plates for imaging. Samples were imaged using a Keyence BZ X710 fluorescence microscope (Osaka, Japan). Z-stacked images were analyzed using Keyence software (BZ Analyzer, Osaka, Japan). ImageJ was used to stitch Figure 2C images together, while all other images were organized using GraphPad Prism.

### AlamarBlue metabolic activity assay

Cell metabolic activity was evaluated using the AlamarBlue assay (ThermoFisher). Culture media was removed from each well and replaced with 200 μL of AlamarBlue reagent solution diluted 1:10 with media. Samples were incubated for 4 h at 37 °C and 5% CO_2_. Following incubation, the fluorescent signal was measured using a plate reader (Thermo Scientific Varioskan Lux, Ex = 555 nm, Em = 595 nm) according to the manufacturer’s instructions. Controls grown in tissue culture plates were seeded at 25,000 cells/well in a 48-well plate, and the media were changed as previously described, with 300 μL per well. Data were analyzed using GraphPad Prism.

### β-Galactosidase assay

β-Galactosidase activity was measured Pierce β-Galactosidase Assay Reagent (Thermo Fisher Scientific) and beta-Galactosidase Assay Stop Solution (Thermo Fisher Scientific) according to the manufacturer’s instructions. Briefly, cells were washed, and then 100 μL of assay buffer was added. Then, the cells were incubated for 30 minutes at 37°C and 5% CO_2_, and 100 μL of stop solution was added before measurement. Absorbance was measured using a plate reader (Thermo Scientific Varioskan Lux) at 405 nm. Data were analyzed using GraphPad Prism.

### LegendPlex Assay

Supernatant was collected from the apical region at each time point after 2 days of cell exposure. Samples were preserved at 20°C until the time the assay was performed. All samples were spun down to remove debris before the assay procedure began. A Legendplex Human Inflammation Panel 1 (BioLegend, Cat# 740808, San Diego, CA, USA) was performed according to the manufacturer’s instructions and run on an Agilent Novocyte Penteon (Agilent, Santa Clara, CA, USA). Files were then analyzed on the BioLegend Legendplex software and graphed in GraphPad Prism. For radar plots, Microsoft Excel and custom Python code were used. Controls for tissue culture plates were performed using a 96-well plate at the same initial seeding density of 25,000 cells/well. The media was changed as previously described, using 300 uL per well.

### Immune score

The immune score was calculated by averaging the replicate values of IL-6, IL-8, and MCP-1 within each condition and day. Data were transformed using log(average + 1) to reduce the influence of differences in absolute magnitude between cytokines. Each cytokine was subsequently standardized across all condition-day groups by calculating z-scores using the formula z = (x−μ)/σ, where μ and σ represent the mean and standard deviation across all groups. The immune score for each condition and day was defined as the arithmetic mean of the z-scores for IL-6, IL-8, and MCP-1. To obtain a single immune score for each condition across the experiment, day-level immune scores were averaged across D2, D7, and D14. The calculations were performed using custom Python code, and the results were graphed in GraphPad Prism.

### Collagen Assay

Media was collected at Days 3, 9, 12, and 14, coinciding with scheduled media changes. At each time point, the full 300 µL of conditioned media was removed from each well prior to replenishment with fresh media. Collected media was used directly for collagen quantification using the Sigma-Aldrich MAK322 fluorometric collagen assay per manufacturer’s protocol using a plate reader (Thermo Scientific Varioskan Lux, Ex = 375 nm, Em = 465 nm). Data were analyzed using GraphPad Prism.

### Immunofluorescence staining

Hydrogels were washed three times with DPBS and fixed with 4% paraformaldehyde (ThermoFisher Scientific) for 20 min at room temperature. Samples were washed with DPBS three times and treated with 0.3% (w/v) Sudan Black B (MilliporeSigma, Burlington, MA, USA) in 70% ethanol for 20 min at room temperature to reduce autofluorescence from silk. Hydrogels were rinsed with DPBS and permeabilized with 0.1% Triton X-100 (Sigma-Aldrich) in DPBS for 1 h at room temperature, followed by DPBS washes. Samples were blocked with 3% FBS in DPBS for 1 h at room temperature. Primary antibody staining was performed overnight at 4°C using anti-α smooth muscle actin (α-SMA) antibody (1:200, mouse monoclonal, Abcam ab7817). All antibody staining was done in diluted DPBS containing 3% FBS. Samples were washed with DPBS and incubated with secondary antibodies for 1.5 h at room temperature using donkey anti-mouse Alexa Fluor 647 (1:500, ThermoFisher Scientific, A-31571). During secondary staining, phalloidin-iFluor 488 (1:500, Abcam, ab176753) was used to label F-actin, and DAPI (1:200, ThermoFisher Scientific, 62248) was used to stain nuclei. Samples were washed with DPBS and imaged. Fluorescence images were acquired using a Leica TCS SP8 Microscope (Wetzlar, Germany) with a 10x/0.3 objective, lasers (DAPI/405 nm, phalloidin/488 nm, Alexa/647 nm), and z-stack images (z-step 2 μm) were collected to visualize cellular organization. Images were processed with Leica Application Suite X (Wetzlar, Germany) software, and settings were maintained consistently across comparable conditions. Brightness/contrast adjustments were applied uniformly during figure preparation. The data was organized in a layout in GraphPad Prism.

### Quantitative PCR (qPCR)

For each experimental condition, four transwells were pooled to generate a single biological sample, and three pooled samples were prepared per treatment group. Total RNA was isolated using the RNA/Protein Purification Plus Kit (Norgen Biotek, Thorold, ON, Canada). Pooled hydrogels (n=4) were placed in 300 µL lysis buffer (Buffer SKP) with β-mercaptoethanol (Fisher Scientific, Waltham, MA, USA) and poly-A carrier RNA (Qiagen #1068337, Hilden, DE, USA), then frozen at -80°C for at least overnight and no more than 7 days. Samples were then thawed, and cells were released from the hydrogels using a Protease XIV (Sigma-Aldrich) digestion step for 30 min at 0.001 U/mL. The resulting lysate was processed for RNA and protein following the manufacturer’s protocol. RNA concentration and purity were characterized via Nanodrop 2000 (ThermoFisher). Contaminating DNA was then removed using the TURBO DNA-free Kit (Invitrogen, Carlsbad, CA, USA). Complementary DNA was generated using iScript (Bio-Rad, Hercules, CA, USA) according to the manufacturer’s protocols. qPCR was performed using Taqman Fast Advanced Master Mix (ThermoFisher Scientific) and the CFX96 Real-Time PCR Detection System (Bio-Rad). Taqman primers can be found in **Table S1**. Gene expression levels of fibrosis-related markers were evaluated relative to the housekeeping gene GAPDH. Relative gene expression was calculated using the comparative Ct (ΔΔCt) method. Ct values for target genes were first normalized to GAPDH to obtain ΔCt values. ΔΔCt values were calculated by comparing each experimental group to the corresponding averaged day 2 initial condition. No-reverse transcriptase (no-RT) controls were included for each sample to confirm the absence of genomic DNA contamination. No amplification was detected in no-RT control reactions, confirming that PCR signals reflect mRNA-derived cDNA. Data were analyzed using Microsoft Excel and GraphPad Prism.

## Declaration of generative AI and AI-assisted technologies in the manuscript preparation process

During the preparation of this work, the authors used Claude (Anthropic) and Grammarly in order to write Python code and edit the manuscript. After using these tools, the authors reviewed and edited the content as needed and take full responsibility for the content of the published article.

## Data Availability Statement

The data that support the findings of this study are available from the corresponding author upon reasonable request.

## Acknowledgments

This work was supported by the US National Institutes of Health under grant number P41EB027062. M.L.A was supported by Award Number K12GM133314 from the National Institute of General Medical Sciences for the Tufts IRACDA Program. In addition, M.L.A. received the Natalie V. Zucker Research Award that supported this work.

## CRediT authorship contribution statement

M.L.A.: Conceptualization, Methodology, Formal Analysis, Investigation, Resources, Writing – Original Draft, Writing – Review & Editing, Visualization, Funding Acquisition. M.S.: Methodology, Investigation, Writing – Review & Editing. A.S.M.: Methodology, Investigation, Writing – Review & Editing. A.Y.: Investigation, Writing – Review & Editing. T.F.: Investigation, Writing – Review & Editing. J.L.D.: Investigation, Writing – Review & Editing. P.L.G.: Investigation, Writing – Review & Editing. S.M.P.: Investigation, Writing – Review & Editing. S.G.: Investigation, Writing – Review & Editing. J.K.S.: Investigation, Writing – Review & Editing. J.J.H.: Investigation, Writing – Review & Editing. G.V.N.: Funding Acquisition, Writing – Review & Editing. D.L.K.: Funding Acquisition, Writing – Review & Editing.

