## Supplemental for "Progressive matrix stiffening of tyramine-modified silk fibroin hydrogels governs stage-specific pulmonary fibroblast activation"

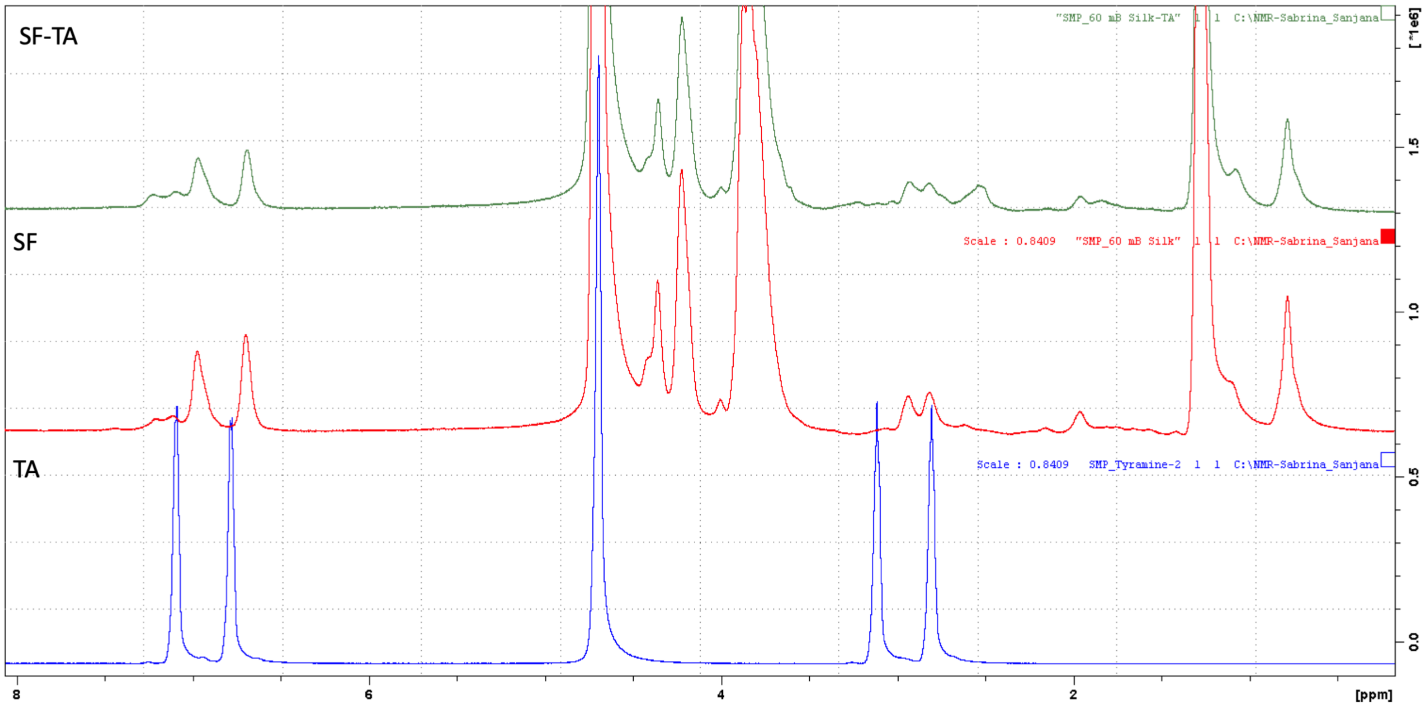


**Supplementary Figure 1: NMR of Tyramine-modified Silk.**

**Supplementary Table 1: TaqMan primers used for qPCR.**

| GENE | CATALOG # | COMPANY |
|---|---|---|
| GAPDH | Hs02786624_g1 | ThermoFisher |
| ACTA2 | Hs00426835_g1 | |
| YAP | Hs00902712_g1 | |
| FAK | Hs01056457_m1 | |
| COL1A1 | Hs00164004_m1 | |
| TGFB1 | Hs00998133_m1 | |
| ITGB1 | Hs01127536_m1 | |
| RHOA | Hs00357608_m1 | |
| MRTFA | Hs01090249_g1 | |
| CTFG | Hs00170014_m1 | |

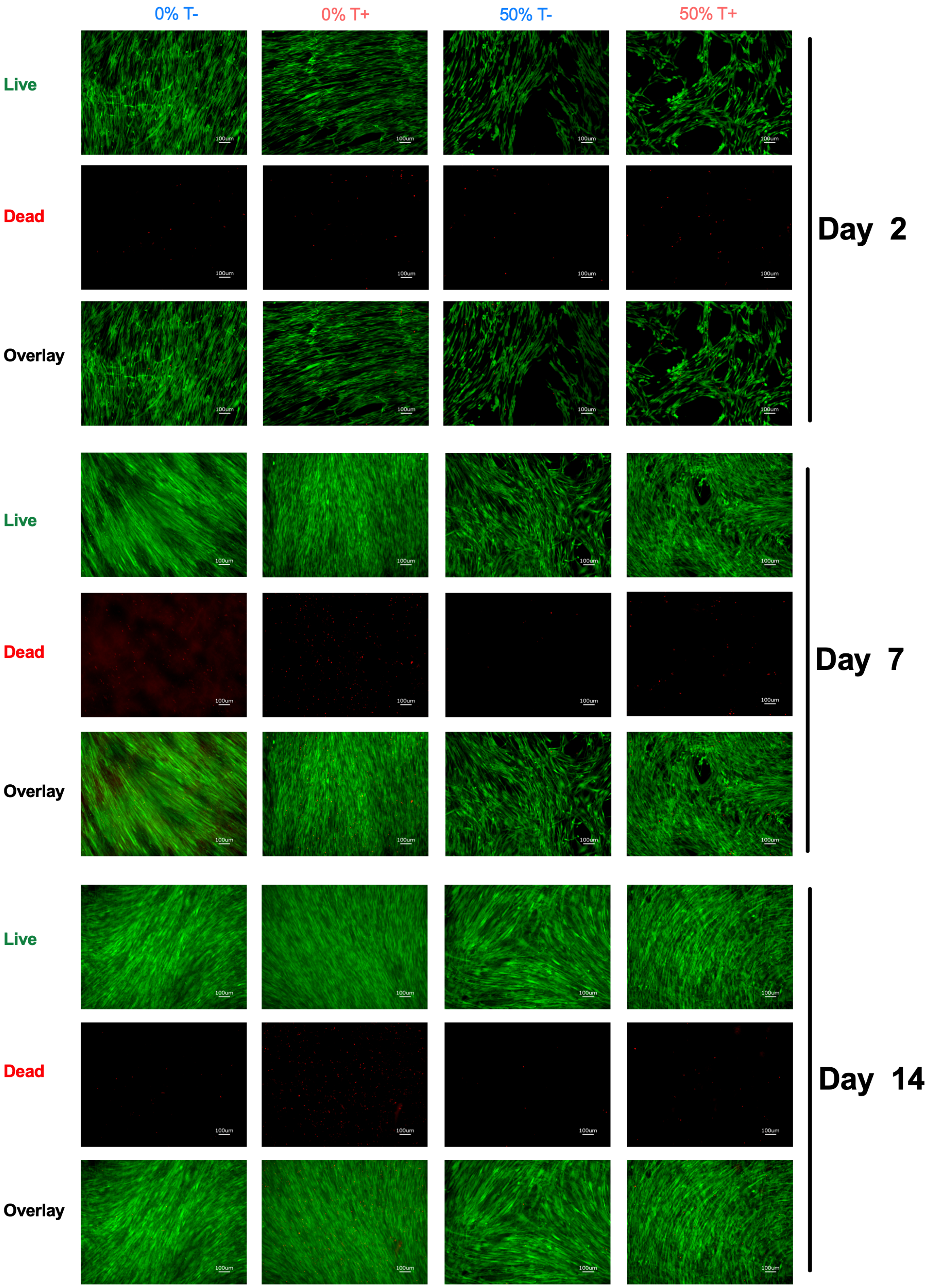


**Supplementary Figure 2: Dynamically stiffening silk hydrogels are cytocompatible over 14 days, biological replicate 1, overlay images part of Figure 2.** NHLFs were seeded at 25,000 cells/well and cultured for 14 days on 0% and 50% SF-TA hydrogels, with or without 5 ng/mL TGFβ. Live/Dead imaging with Calcein AM for live cells (green) and Ethidium Homodimer-1 for dead cells (red) at Day 2, Day 7, and Day 14. Scale bar = 100 μm.

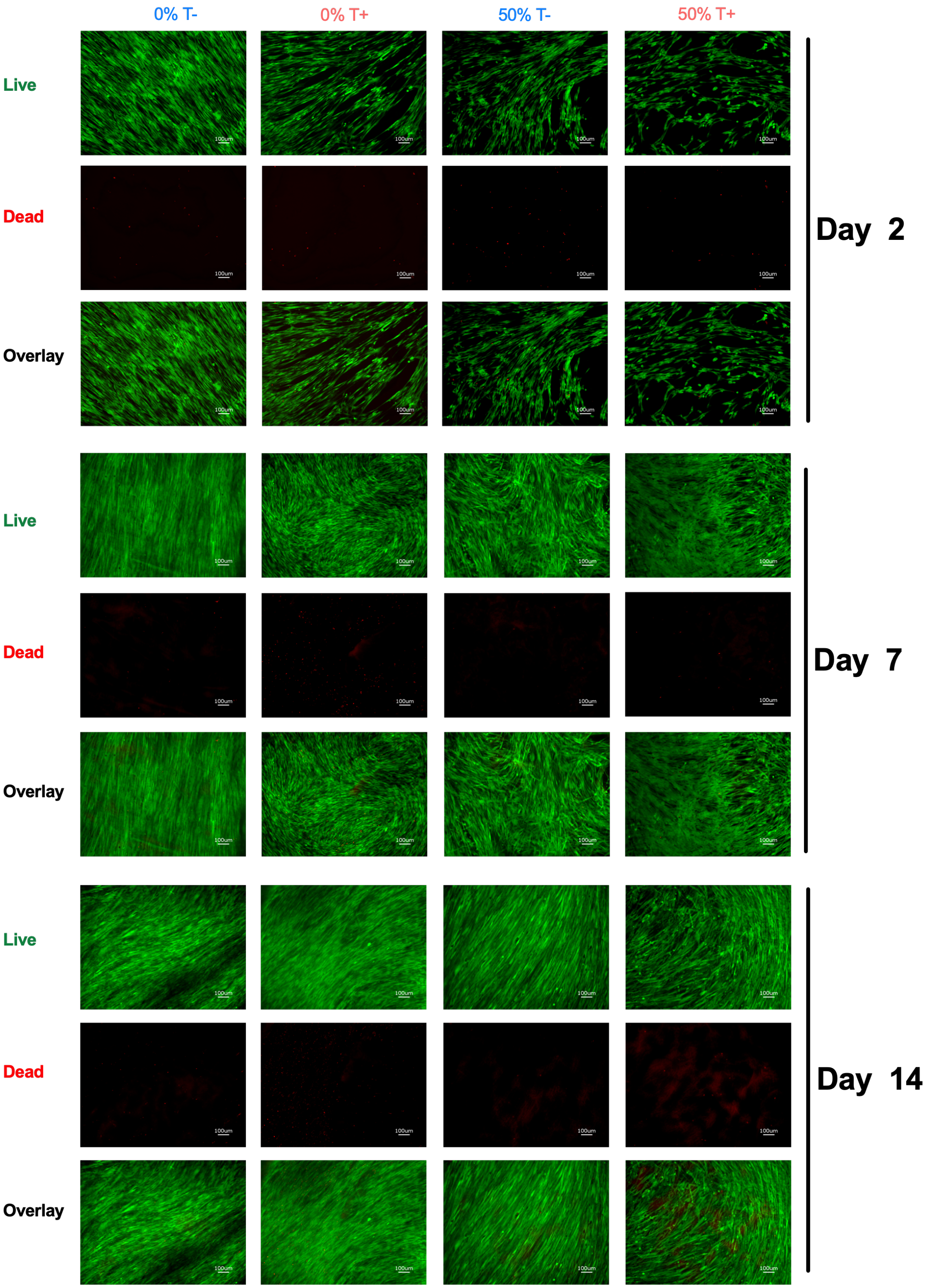


**Supplementary Figure 3: Dynamically stiffening silk hydrogels are cytocompatible over 14 days, biological replicate 2.** NHLFs were seeded at 25,000 cells/well and cultured for 14 days on 0% and 50% SF-TA hydrogels, with or without 5 ng/mL TGFβ. Live/Dead imaging with Calcein AM for live cells (green) and Ethidium Homodimer-1 for dead cells (red) at Day 2, Day 7, and Day 14. Scale bar = 100 μm.

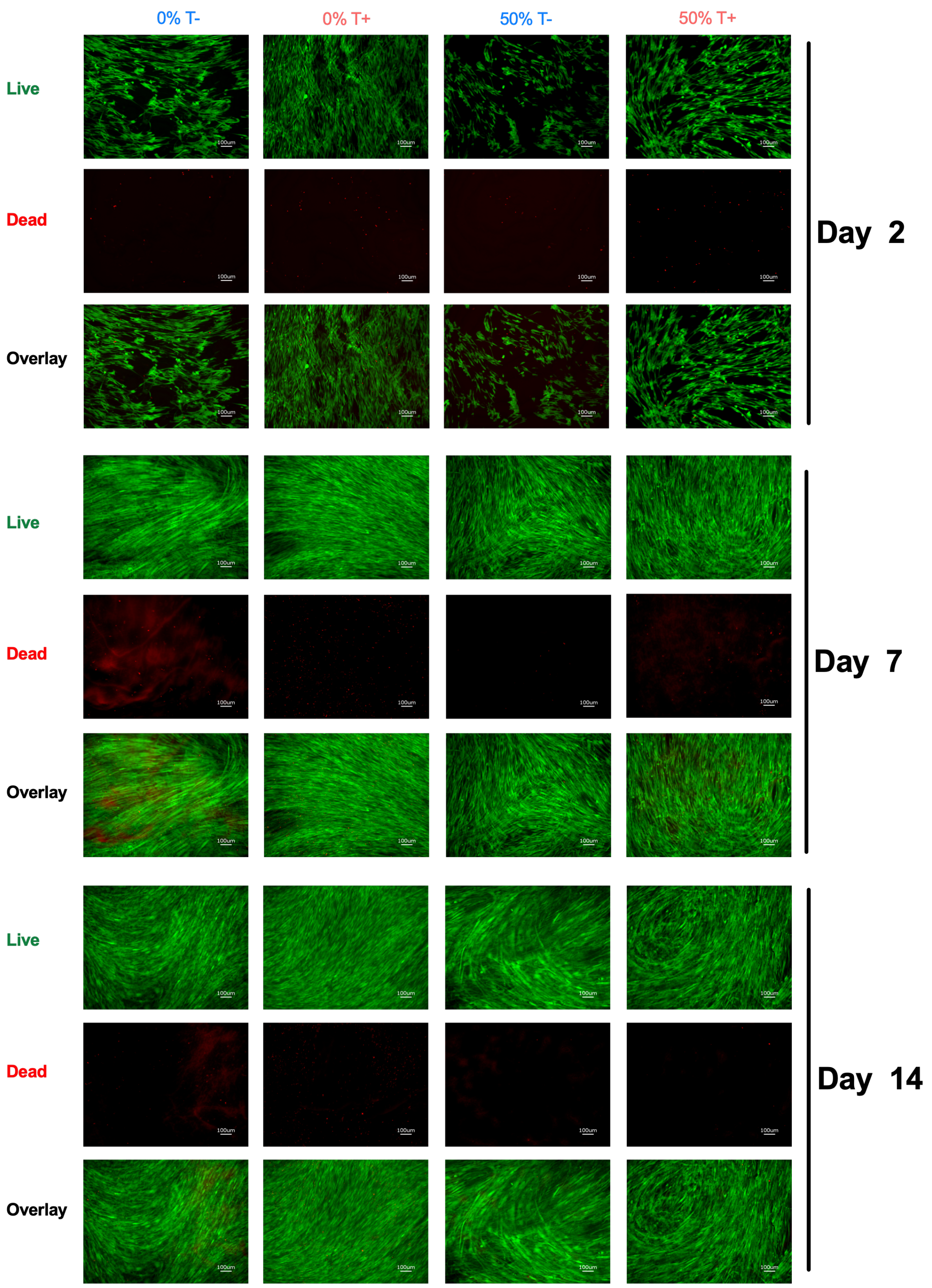


**Supplementary Figure 4: Dynamically stiffening silk hydrogels are cytocompatible over 14 days, biological replicate 3.** NHLFs were seeded at 25,000 cells/well and cultured for 14 days on 0% and 50% SF-TA hydrogels, with or without 5 ng/mL TGFβ. Live/Dead imaging with Calcein AM for live cells (green) and Ethidium Homodimer-1 for dead cells (red) at Day 2, Day 7, and Day 14. Scale bar = 100 µm.

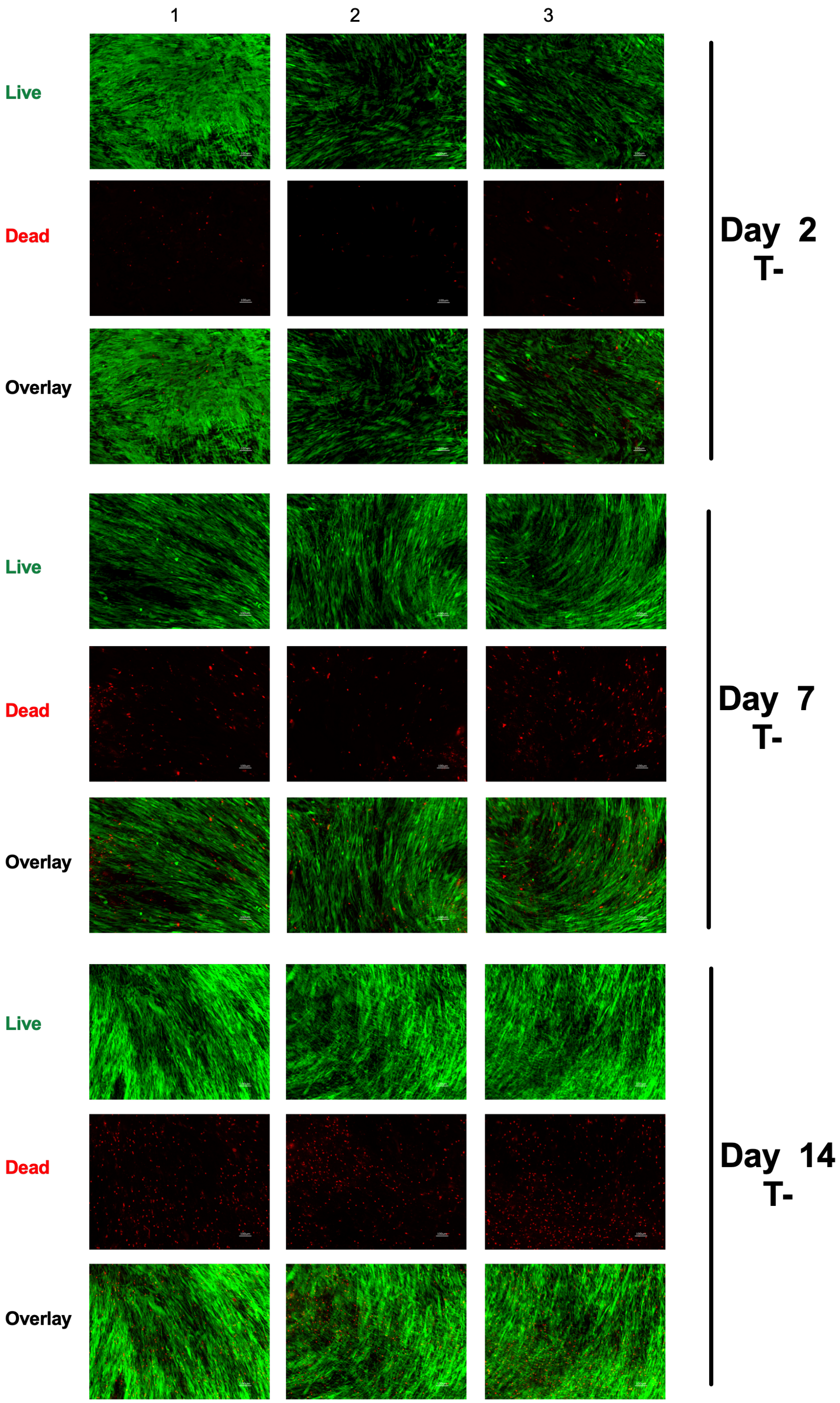


**Supplementary Figure 5: TCP controls for NHLF cells over 14 days, without TGFβ.** NHLFs were seeded at 25,000 cells/well on 96-well tissue culture plastic and allowed to grow for 14 days. Live/Dead imaging with Calcein AM for live cells (green) and Ethidium Homodimer-1 for dead cells (red) at Day 2, Day 7, and Day 14. TCP = Tissue Culture Plate. Scale bar = 100 μm.

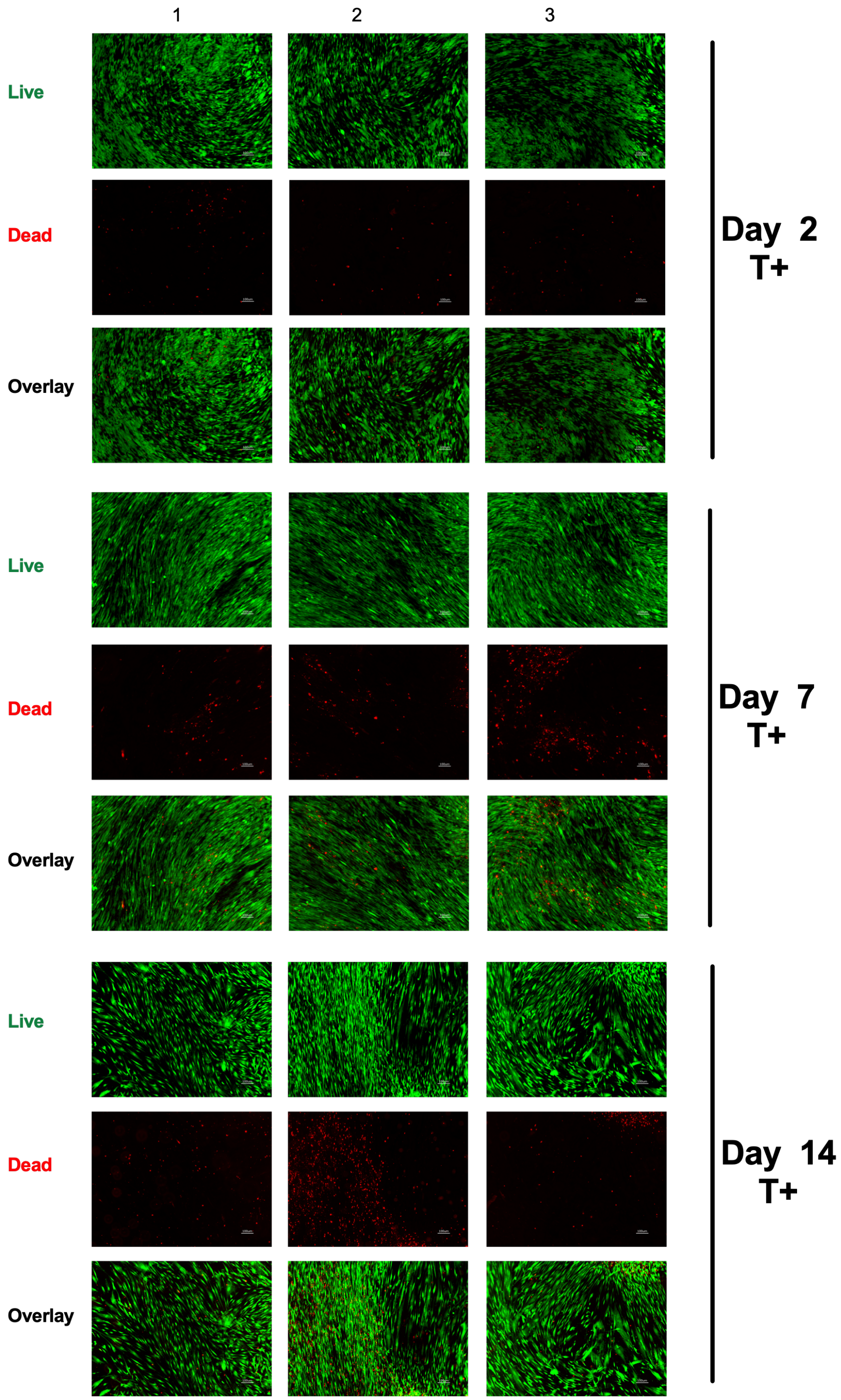


**Supplementary Figure 6: TCP controls for NHLF cells over 14 days, TCP with 5 ng/ml TGFβ.** NHLFs were seeded at 25,000 cells/well on 96-well tissue culture plastic and allowed to grow for 14 days. Live/Dead imaging with Calcein AM for live cells (green) and Ethidium Homodimer-1 for dead cells (red) at Day 2, Day 7, and Day 14. TCP = Tissue Culture Plate. Scale bar = 100 µm.

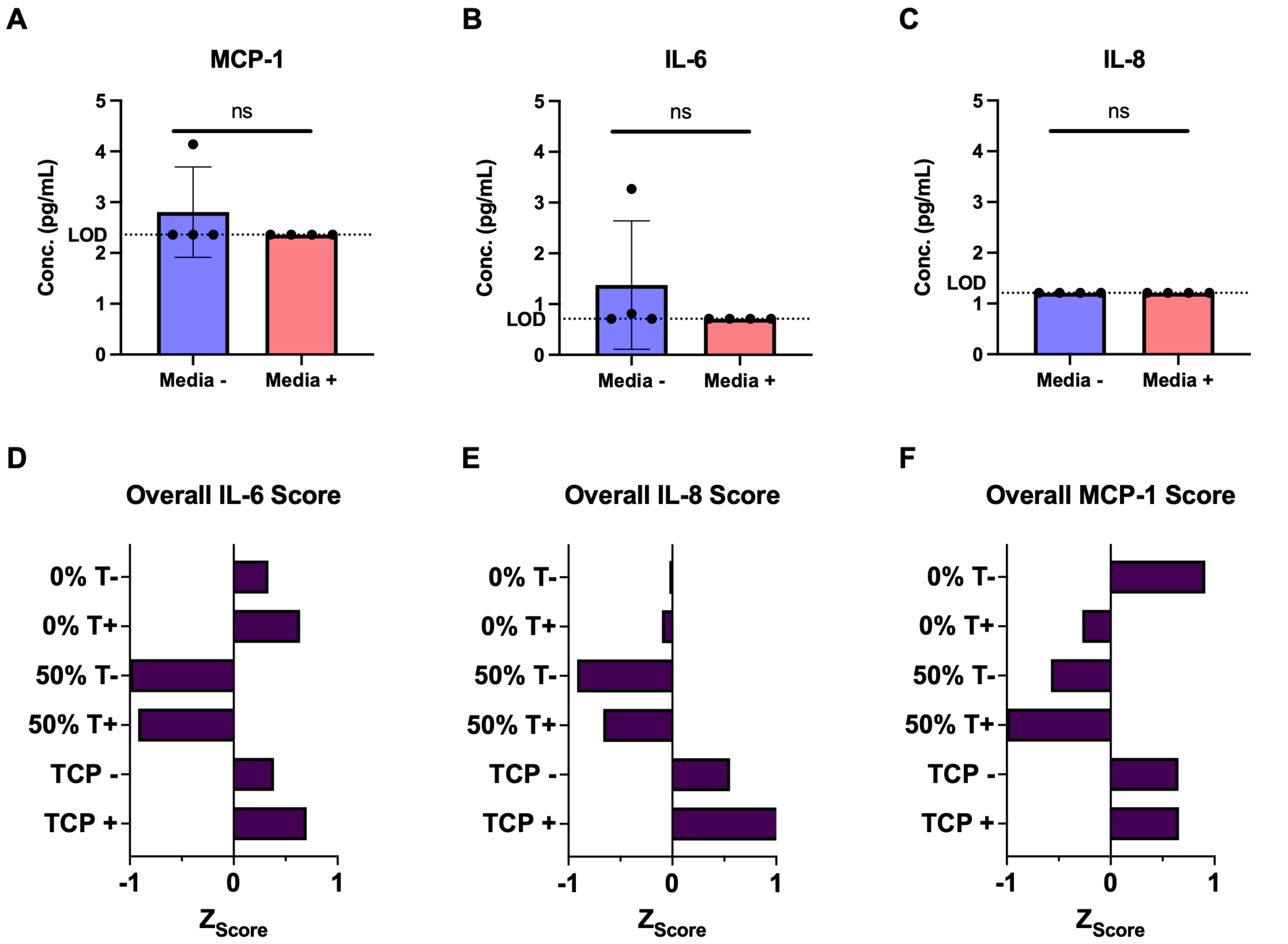


**Supplementary Figure 7: Media controls confirm cytokine measurements are cell-derived.** A-C) Concentrations of MCP-1, IL-6, and IL-8 (pg/mL) in acellular media controls (Media- = T- and Media+= T+) fall at or below the limit of detection (LOD), confirming that detected cytokines in experimental conditions are cell-derived. D-F) Individual Z-scores for IL-6, IL-8, and MCP-1 across all conditions, showing 0% SF-TA and TCP conditions are consistently positive relative to 50% SF-TA. T- = without TGFβ, T+ = 5 ng/mL TGFβ. TCP = Tissue Culture Plate. Welch's t-test (A-C), ns = not significant,

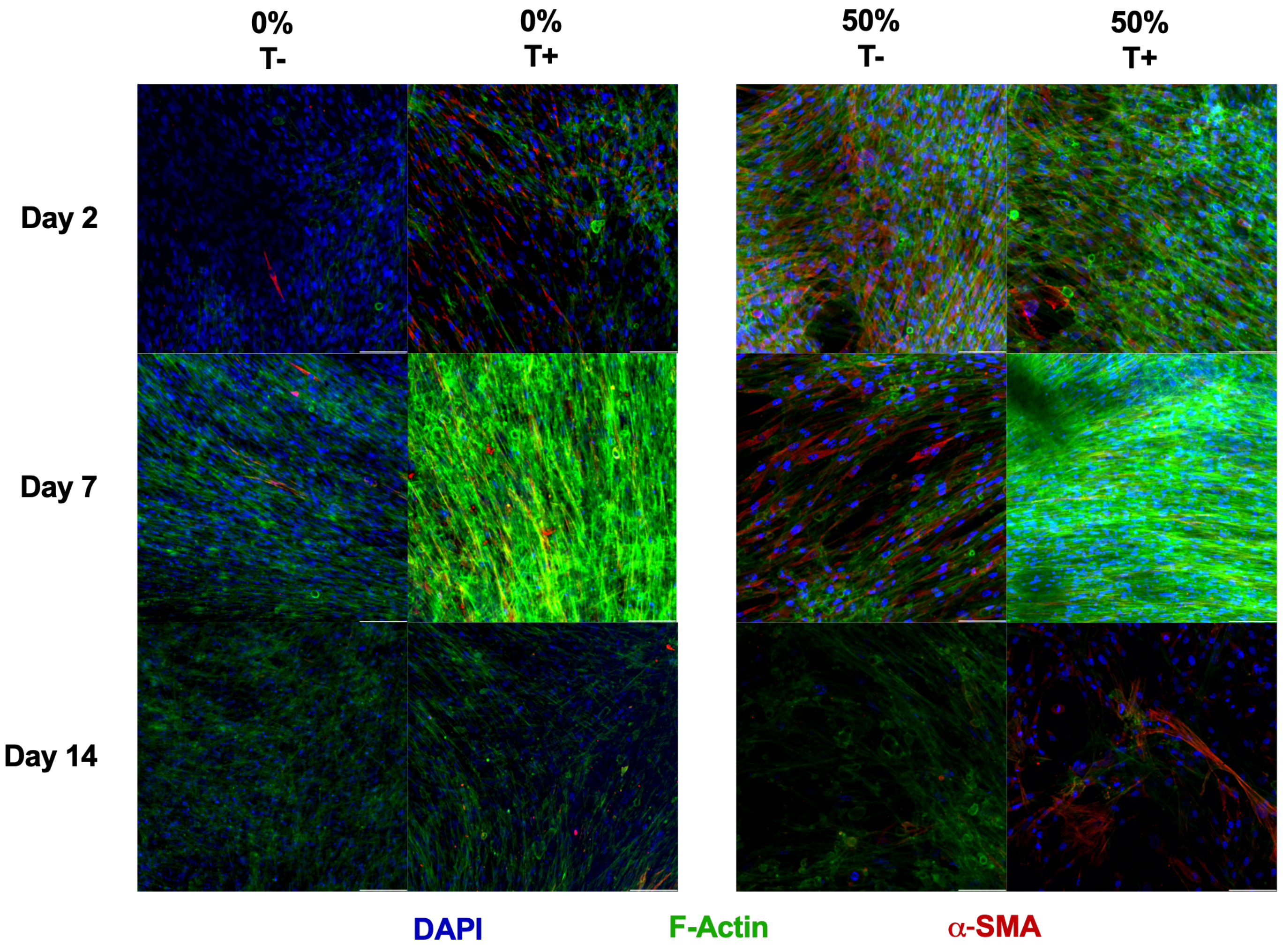


**Supplementary Figure 8: Replicate for immunofluorescence images from Figure 4.** Immunofluorescence images of NHLFs stained for DAPI (blue), F-Actin (green), and α-SMA (red) for 0% SF-TA and 50% SF-TA. T- = without TGFβ, T+ = 5 ng/mL TGFβ. Scale bar = 100 μm.

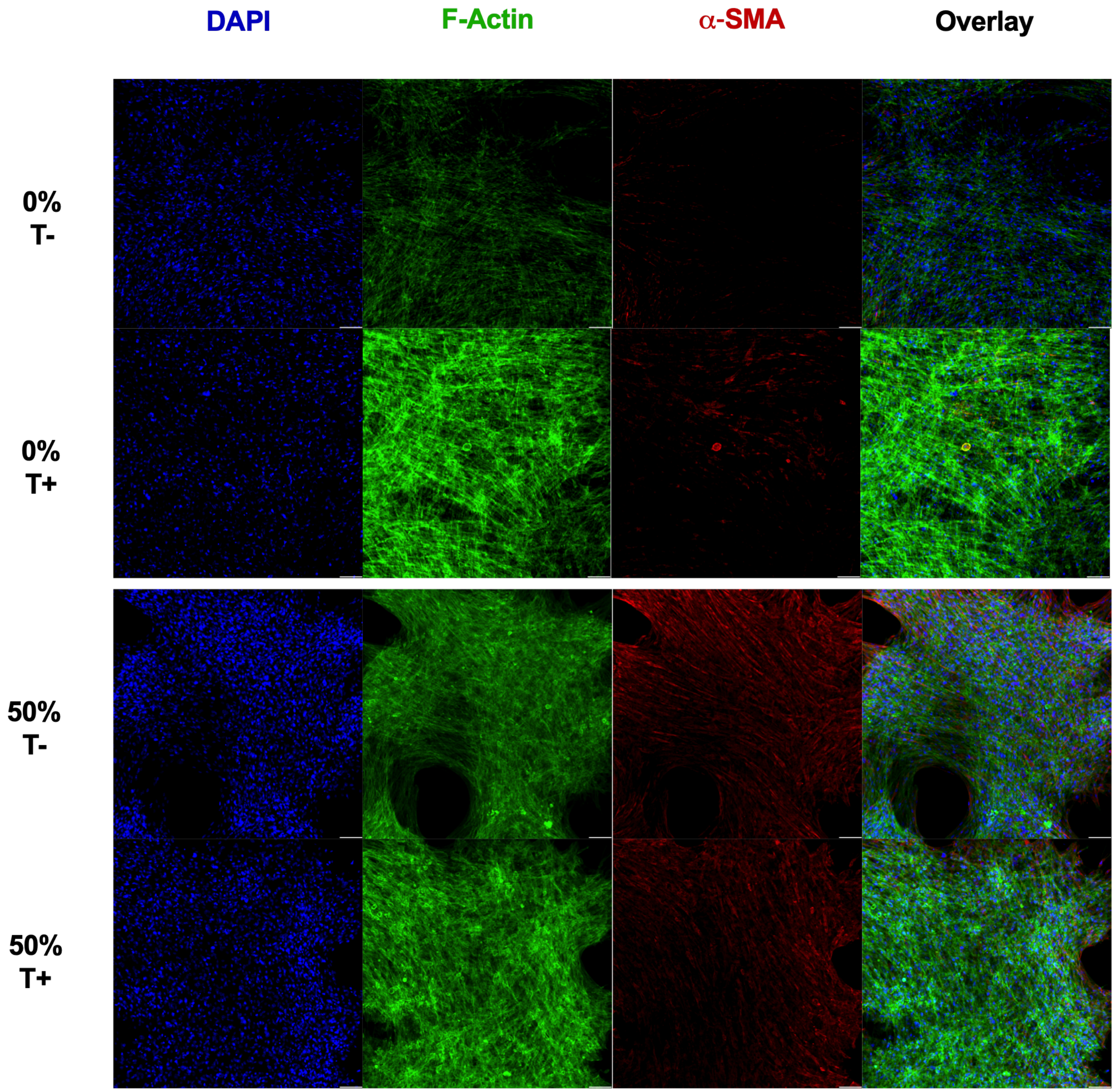


**Supplementary Figure 9: Individual immunofluorescence images from Figure 4 for Day 2.** Immunofluorescence images of NHLFs stained for DAPI (blue), F-Actin (green), and α-SMA (red) for 0% SF-TA and 50% SF-TA. T- = without TGFβ, T+ = 5 ng/mL TGFβ. Scale bar = 100 μm.

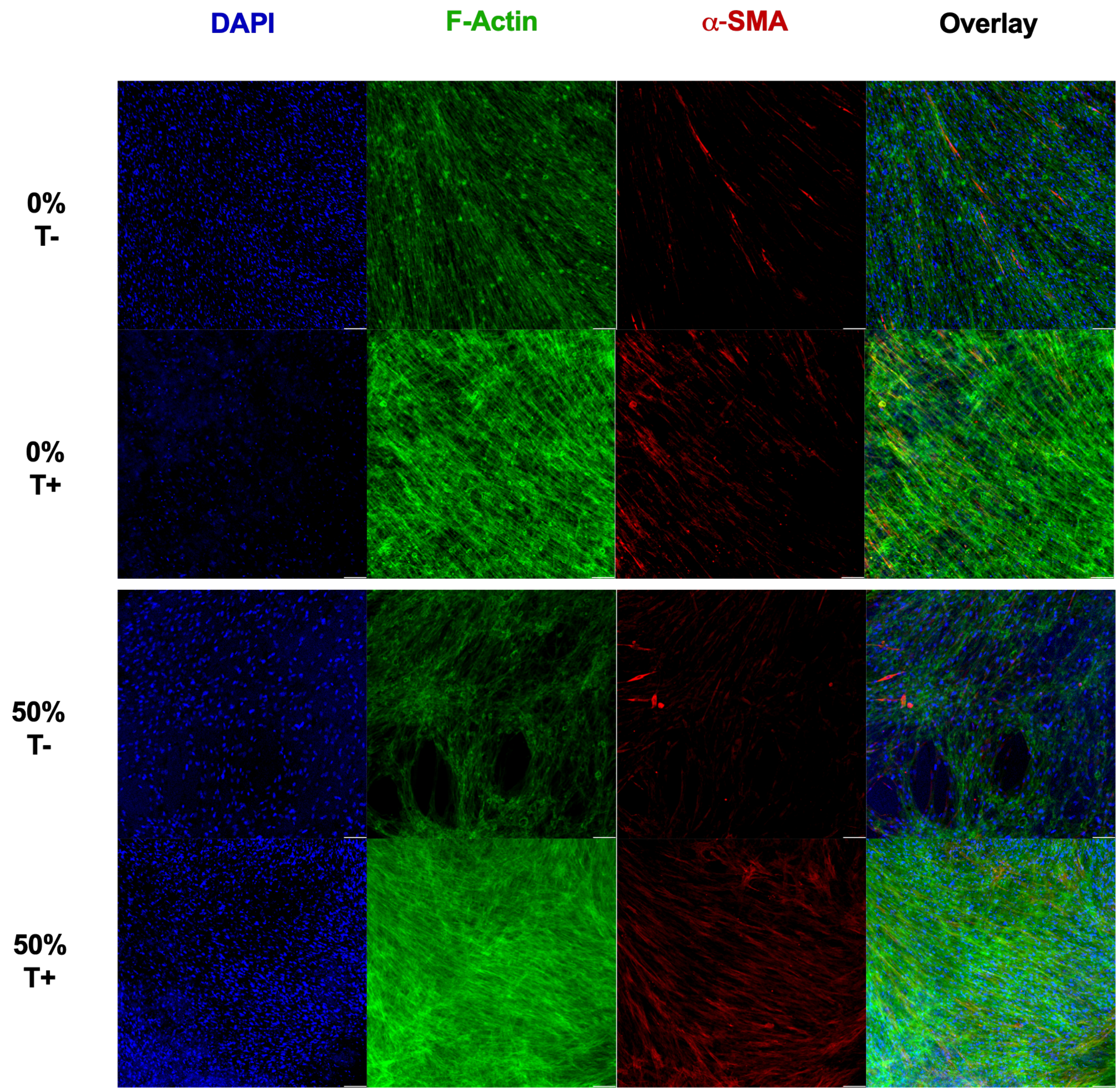


**Supplementary Figure 10: Individual immunofluorescence images from Figure 4 for Day 7.** Immunofluorescence images of NHLFs stained for DAPI (blue), F-Actin (green), and α-SMA (red) for 0% SF-TA and 50% SF-TA. T- = without TGFβ, T+ = 5 ng/mL TGFβ. Scale bar = 100 μm.

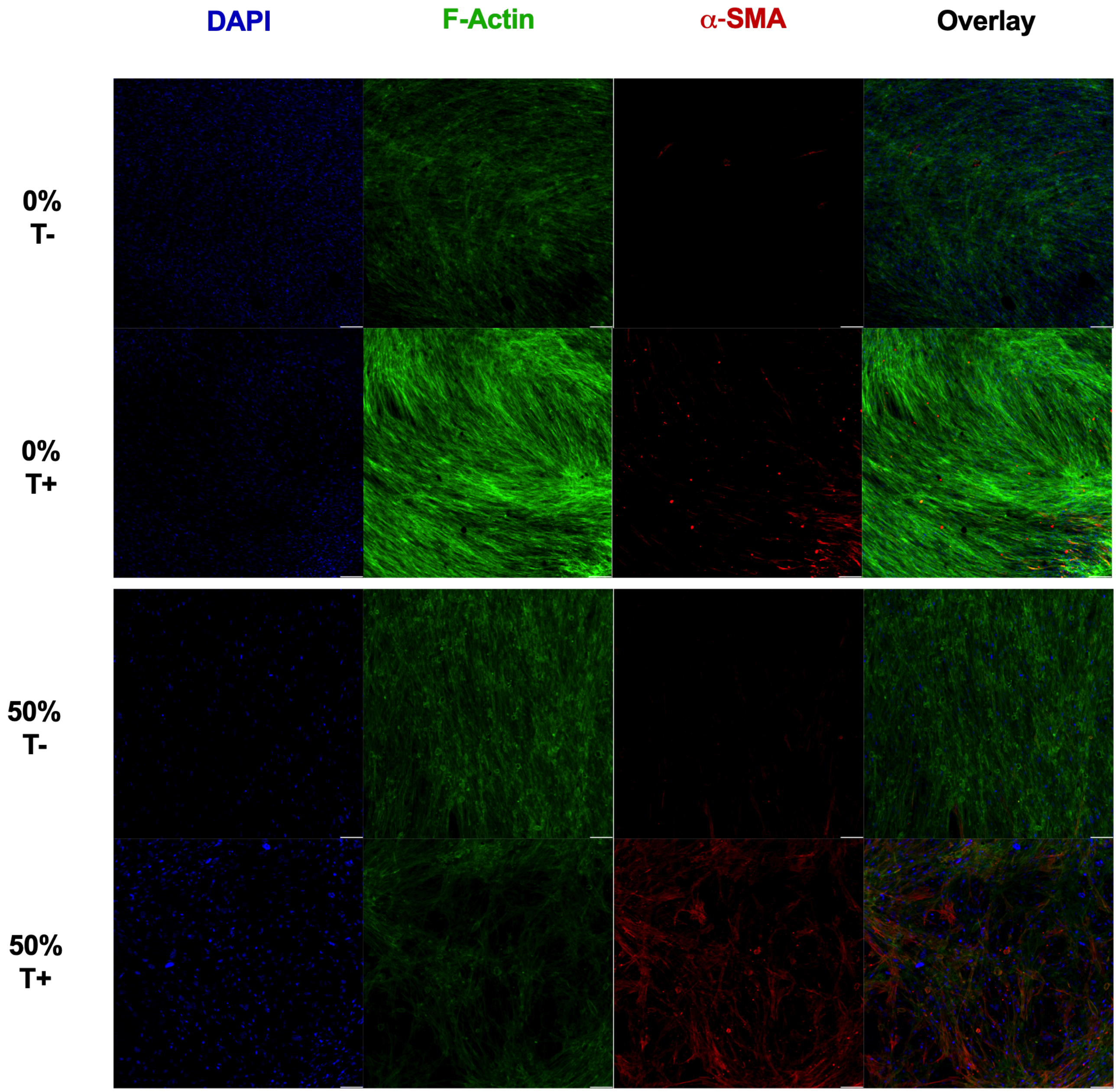


**Supplementary Figure 11: Individual immunofluorescence images from Figure 4 for Day 14.** Immunofluorescence images of NHLFs stained for DAPI (blue), F-Actin (green), and α-SMA (red) for 0% SF-TA and 50% SF-TA. T- = without TGFβ, T+ = 5 ng/mL TGFβ. Scale bar = 100 μm.

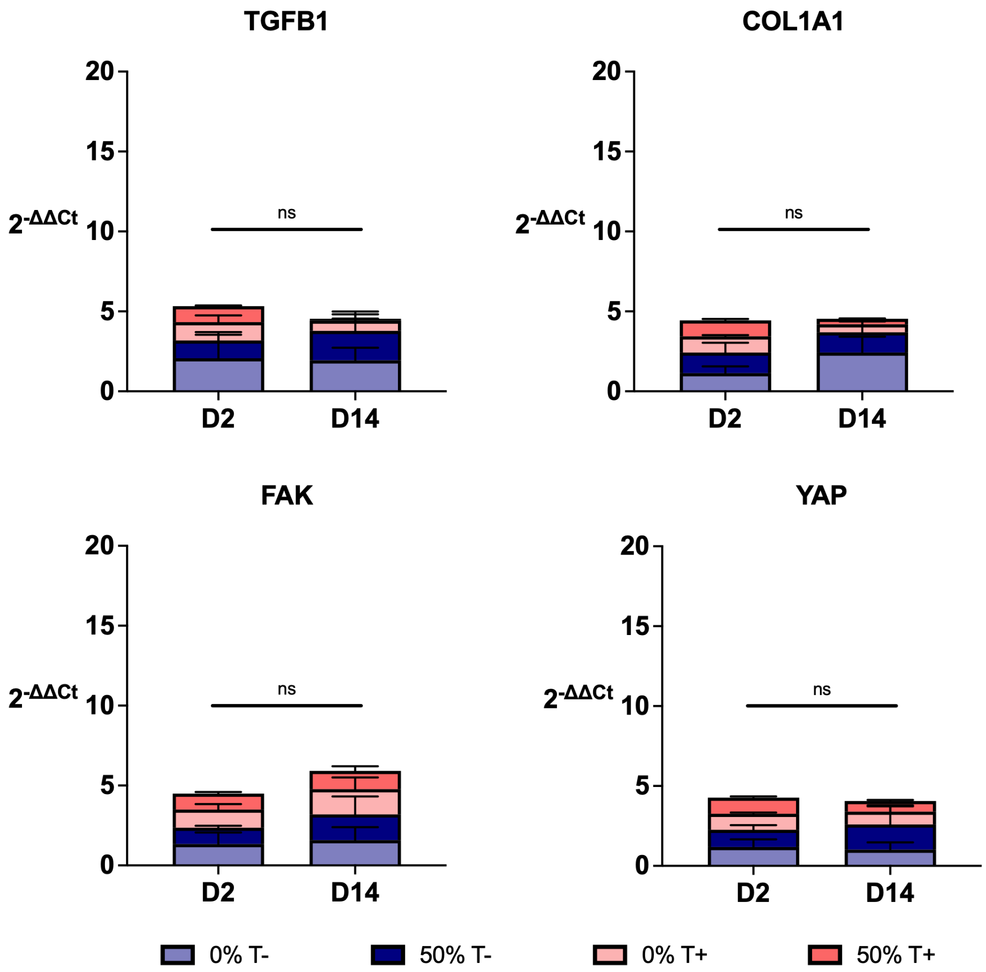


**Supplementary Figure 12: Downstream mechanosensing and fibrotic gene expression markers show no significant change between Day 2 and Day 14.** RT-qPCR expression of TGFB1, COL1A1, FAK, and YAP normalized to GAPDH and to the Day 2 average for each group, reported as $2^{-\Delta\Delta Ct}$ for 0% and 50% SF-TA hydrogels with and without TGFβ at Day 2 and Day 14. T- = without TGFβ, T+ = 5 ng/mL TGFβ. N=3, Error bars = SEM. Welch's t-test, ns = not significant $p > 0.05$.
